# Quantifying the movement, behavior, and environmental context of group-living animals using drones and computer vision

**DOI:** 10.1101/2022.06.30.498251

**Authors:** Benjamin Koger, Adwait Deshpande, Jeffrey T. Kerby, Jacob M. Graving, Blair R. Costelloe, Iain D. Couzin

**Affiliations:** Department of Collective Behavior, Max Planck Institute of Animal Behavior, Konstanz, Germany; Centre for the Advanced Study of Collective Behaviour, University of Konstanz, Konstanz, Germany; Department of Biology, University of Konstanz, Konstanz, Germany; Aarhus Institute of Advanced Studies, Aarhus University, Aarhus, Denmark; Neukom Institute for Computational Science, Dartmouth College, Hanover, USA; Section for Ecoinformatics and Biodiversity, Department of Biology, Aarhus University, Aarhus, Denmark; Advanced Research Technology Unit, Max Planck Institute of Animal Behavior, Konstanz, Germany

**Keywords:** behavioral tracking, computer vision, drones, environmental reconstruction, gelada monkey, pose, posture, video analysis, wildlife, zebra

## Abstract

1. Methods for collecting animal behavior data in natural environments, such as direct observation and bio-logging, are typically limited in spatiotemporal resolution, the number of animals that can be observed, and information about animals’ social and physical environments.
2. Video imagery can capture rich information about animals and their environments, but image-based approaches are often impractical due to the challenges of processing large and complex multi-image datasets and transforming resulting data, such as animals’ locations, into geographic coordinates.
3. We demonstrate a new system for studying behavior in the wild that uses drone-recorded videos and computer vision approaches to automatically track the location and body posture of free-roaming animals in georeferenced coordinates with high spatiotemporal resolution embedded in contemporaneous 3D landscape models of the surrounding area.
4. We provide two worked examples in which we apply this approach to videos of gelada monkeys and multiple species of group-living African ungulates. We demonstrate how to track multiple animals simultaneously, classify individuals by species and age-sex class, estimate individuals’ body postures (poses), and extract environmental features, including topography of the landscape and animal trails.
5. By quantifying animal movement and posture, while simultaneously reconstructing a detailed 3D model of the landscape, our approach opens the door to studying the sensory ecology and decision-making of animals within their natural physical and social environments.

## Introduction

Studying animals in the wild is essential for understanding how environments shape behavior, and how behavior affects individual fitness. The impacts of animals’ behaviors also extend beyond the individual to broader ecological scales: the cumulative behavior of many interacting individuals underpins emergent group-level behaviors such as collective sensing (Berdahl et al., 2013), migration (Flack et al., 2018), and self-assembly (Lutz et al., 2021); mediates population growth and decline (Bro-Jørgensen et al., 2019); and drives ecological processes such as seed dispersal, pollination, and trophic cascades (Atkins et al., 2019; Costa-Pereira et al., 2022). While conservation initiatives commonly focus on preserving phenomena at these broader scales (Bro-Jørgensen et al., 2019) a mechanistic understanding of the individual-level animal behaviors that drive such phenomena is often critical to the success of conservation efforts, including habitat restoration (Hale et al., 2020), wildlife translocation (Berger-Tal et al., 2020), preservation of ecosystem services (Mortelliti, 2022), and human-wildlife conflict interventions (Pekarsky et al., 2021). Thus, the on-going biodiversity crisis (Ceballos et al., 2020) has created an urgent need to understand the drivers and diversity of animal behavioral patterns and how these are influenced by anthropogenic impacts (Buchholz et al., 2019).

Connecting individual behavior to broader scale phenomena requires detailed data on the behavior of large numbers of animals, as well as their interactions with social partners and their biotic and abiotic environments (Costa-Pereira et al., 2022). For example, understanding the collective movement decisions of animal groups requires data on individual movements and how these relate to the concurrent actions of social partners and salient environmental features (Strandburg-Peshkin et al., 2017). Likewise, predicting the resilience of animal-mediated seed dispersal to environmental change requires knowledge of the foraging preferences and movement patterns of dispersers and how these behaviors vary between individuals and across environmental conditions (Mortelliti, 2022; Pollux, 2017). The challenge of collecting such detailed and multifaceted datasets constitutes a major barrier to integrating behavioral ecology – and its individual-level focus – into the broader fields of ecology and conservation, where practitioners primarily rely on species or populations as the units of behavioral analysis (Bro-Jørgensen et al., 2019; Costa-Pereira et al., 2022). The rapid development of animal-borne data-logging devices (known as bio-loggers) in recent decades has transformed our ability to collect high-resolution data on individual animals (Kays et al., 2015), and has fueled calls to incorporate within-species variation into conservation decisions and investigations of ecological processes (Bro-Jørgensen et al., 2019; Mortelliti, 2022; Shaw, 2020). However, beyond the normal constraints of biologging (Hughey et al., 2018), integrating bio-logging data from individual animals with the concurrent social and ecological context remains a significant challenge (Costa-Pereira et al., 2022; Hughey et al., 2018).

Aerial video-based observation, in combination with machine learning-based image processing tools, is emerging as a promising approach for generating the high-resolution movement datasets necessary for quantitative, multi-scale studies of wildlife behavior (Hughey et al., 2018; Tuia et al., 2022). Image-based tracking methods, reviewed in Dell et al. (2014) in which the location of multiple individuals (Walter & Couzin, 2021) and possibly individual body parts (Graving et al., 2019; Mathis et al., 2018; Nath et al., 2019; Pereira et al., 2020) are tracked over the course of a video observation, originated in laboratory settings where overhead cameras enabled continuous monitoring with minimal visual occlusions. Crucially, controlled recording conditions enabled the use of automated tracking solutions making large-scale video-based studies tractable. In recent years, attempts have been made to replicate this approach in field settings, often by using drones to collect aerial imagery of study subjects. Initially, complex visual environments and a lack of controlled lighting and filming conditions prevented automated image-based tracking, leading researchers to rely on alternative approaches, such as manually extracting data from videos (Sprogis et al., 2020) analyzing a subset of still frames (Inoue et al., 2019) or approximating animal movement paths using the drone’s position (Raoult et al., 2018). However, recent developments in deep learning based approaches (Zhao et al., 2019), have simplified the automatic detection of objects in complex scenes, enabling detection of animals and possibly tracking of their movement and body posture in imagery captured with drones (Corcoran et al., 2021) as well as camera traps, smartphones, and thermal, infrared and conventional cameras (Tuia et al., 2022).

An unresolved challenge is the projection of animal locations throughout an observation from the pixel coordinates of the camera to geographic coordinates and standard units of distance (Haalck et al., 2020), a task that is complicated by the complex topography of natural landscapes and the movement of drone-mounted cameras. Multi-camera 3D imaging can surmount these issues, but such systems are logistically and computationally difficult to deploy (Francisco et al., 2020). Instead, previous drone-based behavioral studies have estimated distances using animal body-lengths and approximated the ground as locally flat (Ringhofer et al., 2020; Torney et al., 2018). While appropriate in some landscapes, these approaches are not generalizable and preclude fusion of animal movement data with external georeferenced datasets and analysis of data in standard units.

While recording focal animals, cameras simultaneously capture imagery of the surrounding environment, which can potentially be processed to quantify habitat structure, detect landscape features, or assess resource quality. Previous studies have manually extracted concurrent behavioral and environmental information from imagery captured with camera traps (Hofmeester et al., 2020; C. Sun et al., 2021) and animal-borne cameras (Thompson et al., 2015). But, while aerial photography is widely used for environmental monitoring (Manfreda et al., 2018), to our knowledge, the simultaneous recording of the environment and animal behavior remains unexplored with overhead filming methods.

Outside of concurrent imagery, current methods for integrating movement and behavior with environmental data involve fusing datasets from different sources (e.g. animal location data from biologgers and satellite imagery), or using secondary sensors (such as animal-borne cameras) mounted on biologging tags alongside geolocators (Williams et al., 2020). But these approaches are limited by mismatches between the scale and capture time of different data types (in the case of data fusion), or limited sampling range of sensors (in the case of animal-borne cameras, which have limited fields of view and are prone to obstruction) (Costa-Pereira et al., 2022; Naganuma et al., 2021). Processing video-based observations to generate concurrent behavioral and environmental datasets at consistent spatiotemporal scales could facilitate novel studies on the behavioral interactions of animals with their social, biotic and abiotic environments, thereby generating a more holistic understanding of the individual-level drivers of higher-order ecological processes and patterns (Costa-Pereira et al., 2022).

Here we describe a method for using aerial video and computer vision to collect high-resolution georeferenced locational and behavioral data on free-ranging animals without capturing or tagging them. In our approach, we use drones to record overhead video of focal animals and subsequently use a deep learning-based pipeline to automatically locate and track all individuals in the video. We use Structure-from-Motion (SfM) techniques to reconstruct the 3D topography of the surrounding habitat (D’Urban Jackson et al., 2020) which, combined with further image processing, can accurately transform animal movement data into geographic coordinate systems independent of camera movement or landscape smoothness. We simultaneously generate additional behavioral and environmental information, including estimates of each animal’s body posture (pose) and landscape features such as animal trails. This method, illustrated in Fig. 1, is applicable to a wide range of species, generates behavioral data at sub-second and sub-meter resolution, and produces synchronous information about the surrounding physical and biotic environment. The use of inexpensive consumer drones that can be flexibly redeployed promotes larger sample sizes and replication of studies without incurring additional equipment costs. Although not without its own limitations, such as possible animal disturbance by drones and limits on observation time (see *Limitations and Considerations*), this approach has many advantages that make it a powerful complement to existing methodologies. Our approach thus has the potential to broaden the scope of behavioral ecology to encompass questions and systems that are unsuitable for direct observation or bio-logging approaches in isolation.

**Figure 1.**
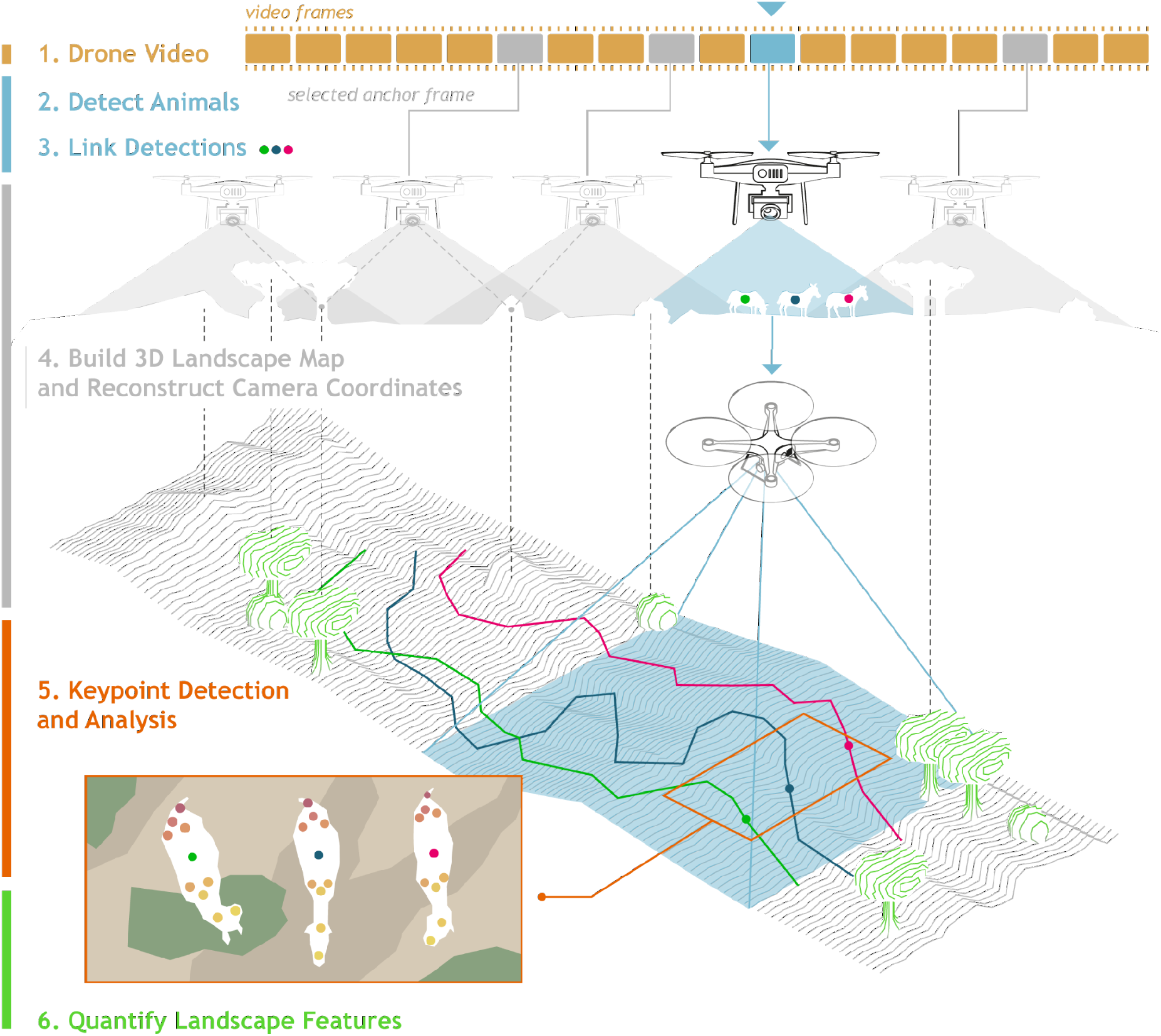
Overview of the processing pipeline for extracting movement, behavioral, and landscape data from aerial drone footage of wildlife. Numbered steps correspond to the numbered sections in the text. First, the animals of interest are video recorded from above (Step 1). Next, an object detection algorithm is used to localize each animal in every video frame (Step 2), and these locations are then linked across frames to generate movement trajectories in pixel coordinates (Step 3). In parallel, anchor frames are selected from the footage and used to build a 3D model of the landscape and estimate the locations of the drone across the observation. Camera locations at anchor frames and local visual features are combined to estimate camera locations for all frames allowing the transformation of the animal trajectories from Step 3 into geographic coordinate space (Step 4). Optionally, further analyses can be performed to extract more detailed behavioral and landscape information, for example through localization of specific body parts (keypoints) for each individual (Step 5) and landscape feature detection (Step 6).

## Methods

Below we outline the major steps of our methodological approach (Fig. 1). Full details of each step are given in the Supplement. To illustrate our processing pipeline, we provide two worked examples with complete code and data available on GitHub (https://github.com/benkoger/overhead-video-worked-examples). We encourage readers to explore these examples and to modify the code for their own datasets.

In the first example, we apply our method to multiple African ungulate species. We recorded ungulate groups at Ol Pejeta and Mpala Conservancies in Laikipia, Kenya over two field seasons, from November 2 to 16, 2017 and from March 30 to April 19, 2018. In total, we recorded thirteen species, but in our example we analyze a single observation of the endangered Grevy’s zebra (*Equus grevyi*) (Rubenstein et al., 2016). We flew DJI Phantom 4 Pro drones [DJI, Shenzhen, China] at an altitude of 80m above the takeoff point. We deployed two drones sequentially in overlapping relays to achieve a continuous 50 minute observation. Drones were piloted manually, and the drone’s position was adjusted as necessary to keep all herd members in frame. While the zebras seemed undisturbed in the example observation and in most other observations, in cases where the animals appeared agitated or fled (as a result of detecting the drone, or our presence via some other cue(s)), we ended the observation. Our method is agnostic to the exact drone-type used, and is scalable to the higher-resolution video made possible by employing more recent models.

In the second worked example, we process video recordings of grassland-dwelling gelada monkeys (*Theropithecus gelada*). Aerial video recordings of gelada monkeys were provided by the Guassa Gelada Research Project. The recordings were collected between October 16, 2019 and February 28, 2020 at the Guassa Community Conservation Area in the Ethiopian highlands. Drones were piloted manually, taking off initially several hundred meters away from the herd and ascending to 100m above ground level before slowly moving above the herd and descending to 70m. This study herd was habituated to close-proximity research observers and construction machinery from nearby roadwork, and appeared undisturbed by the drone apart from an initial increase in glance rate which decreased with exposure. Flights lasted between 5 and 20 minutes. Geladas were recorded with a DJI Mavic 2 Pro [DJI, Shenzhen, China].

In each worked example, we generate movement trajectories for each animal in the example videos and 3D models of the surrounding landscape. In the ungulate example, we also track body keypoints and analyze landscape imagery to detect animal trails, which zebras may follow while moving across the landscape. For the geladas, we train our detection model (Step 2), to distinguish between adult males and other individuals.

### Step 1. Video Recording

Videos should be recorded from above the animals of interest with the camera pointing directly down (nadir view). The visibility of the animals is of vital importance and will be affected by the habitat type, video resolution and characteristics of the animals themselves. As a general rule, videos in which the animals are easily detectable by humans will significantly ease processing. Although many camera platforms can be used to capture the footage, drone-mounted cameras will likely be the most common approach and subsequently we use “drone” and “camera” interchangeably. While our general approach may work for fixed wing drones that are constantly in motion, this method was designed for copter style drones that can transition between periods of hovering and movement. For specific points to consider when planning drone-based data collection, see *Limitations and Considerations*, below and, for more details, *Step 1* in the Supplement.

### Step 2. Detection

Animal detection and localization in each video frame is accomplished with deep convolutional neural networks (CNNs), which we build, train and deploy to predict localizing **bounding boxes** (see Table 1 for definitions of bolded terms) for all individuals in all frames of each video. We first manually annotate frames from the video footage to build image sets for training the model and evaluating its performance. **Annotation** can be tedious, and the content of the images will strongly affect model performance; therefore, it is important to carefully consider the best annotation strategy to achieve high information value for each annotation while minimizing human labor (see Supplement section 2.1).

**Table 1.**
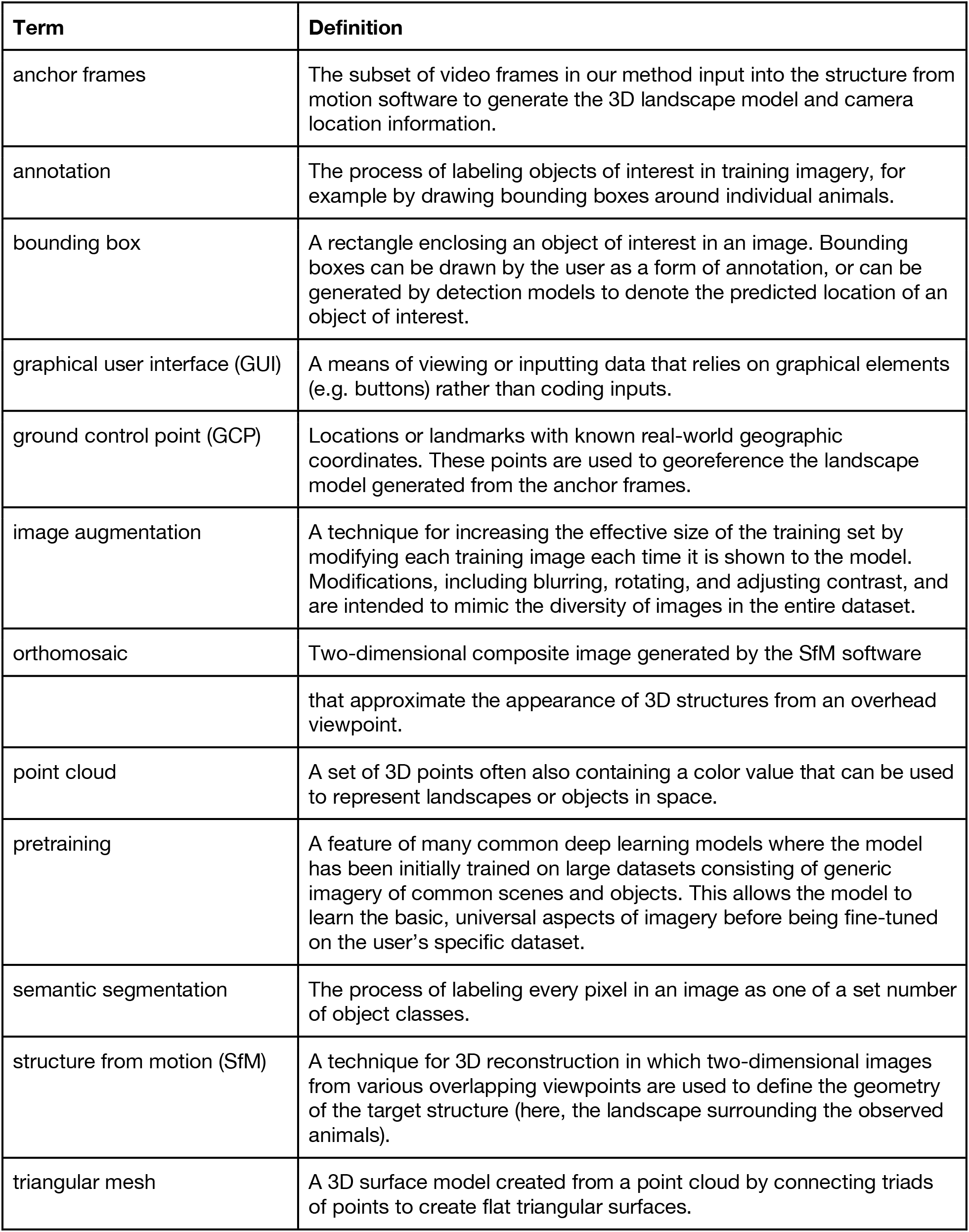
Glossary of terms. Defined terms are bolded at first appearance in the text.

To train a deep learning-based object detection model, researchers must first choose an appropriate software framework and a specific model to train within that framework (see Supplement section 2.2). We use the Detectron2 API within the PyTorch framework (Paszke et al., 2019; Wu et al., 2019) in our worked examples, but the user may choose a different framework depending on their level of coding proficiency or prior experience with other programming libraries. For simple use cases with clearly-visible individuals, many common models can be readily re-configured for the researchers’ data. Users with more challenging footage may need to make more considered choices and possibly incorporate more recent high-performance algorithms. Generally, users should choose models that have been **pretrained** on general image datasets and that incorporate **image augmentation**, as these steps increase training efficiency and require smaller annotated training sets. See Table S1 for annotation statistics and model performance metrics for the datasets and trained models used in the worked examples.

After the model is trained, all frames from all videos can be automatically processed. For each video frame the model generates bounding box coordinates, predicted object classes and confidence scores for every detected object (Fig. 2). We initially take the mean of the coordinates of the bounding box corners as an individual’s location in the frame, which we subsequently use for tracking.

**Figure 2.**
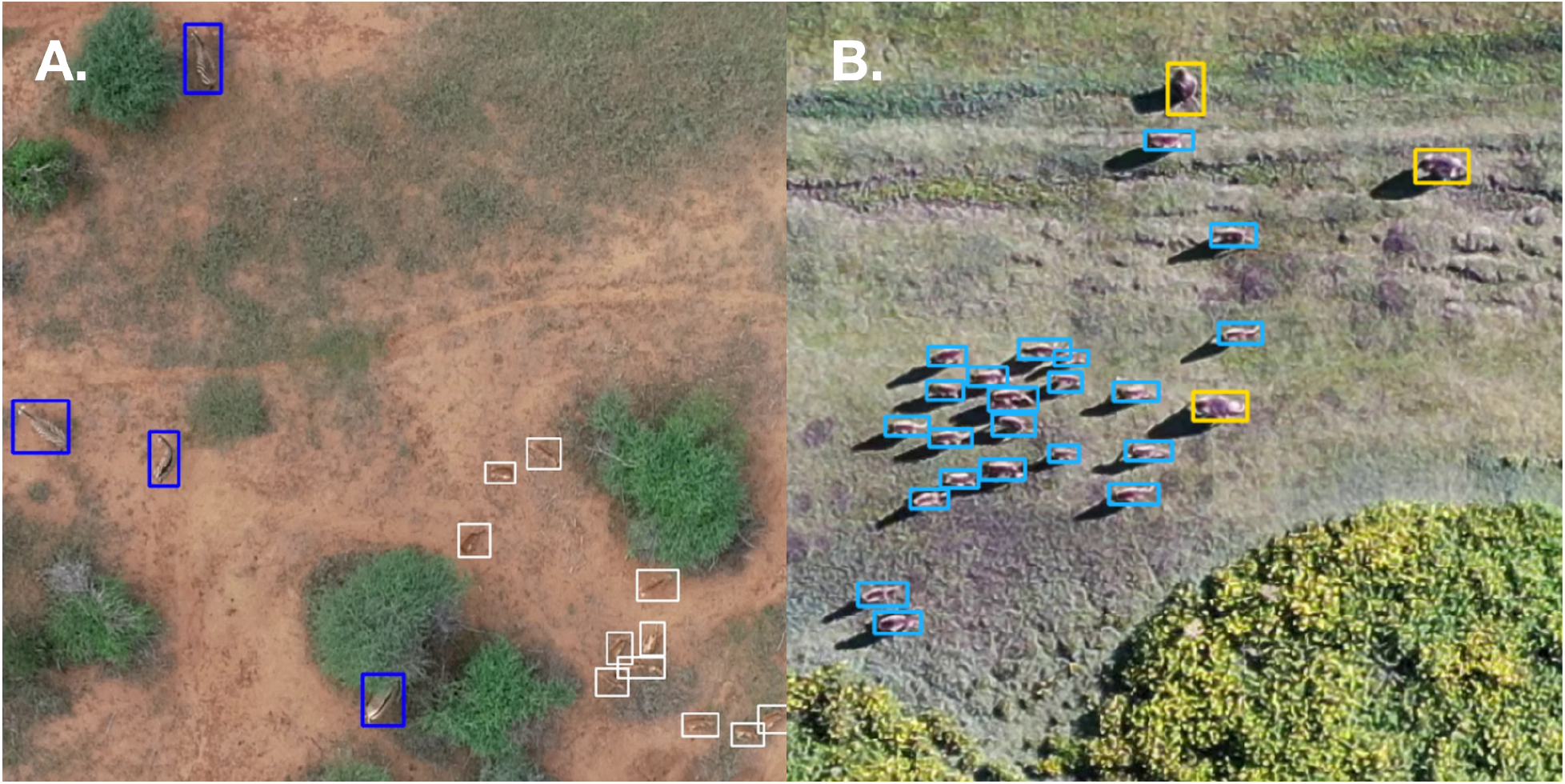
Predicted bounding boxes for two example video frames using the trained models from our worked examples. A) In the ungulates example, the model distinguishes between five classes (zebra, impala, buffalo, waterbuck, and other). The animals in dark blue bounding boxes are Grevy’s (*Equus grevyi*) and plains zebras (*E. burchelli*) and animals in white bounding boxes are impala (*Aepyceros melampus*). B) In the gelada monkey example, we distinguish between species and age-sex class (human-observer, adult-male gelada, and other-gelada). Yellow boxes are predicted adult-males while light blue boxes are a mix of females and juveniles.

### Step 3. Tracking

Tracking, or linking positions across video frames, allows us to generate trajectories for all detected individuals in the pixel coordinate system of the video. To match individual locations across consecutive frames, we use a modified version of the Hungarian algorithm (Kuhn, 1955), which finds the pairing of trajectories and new positions that minimizes the total distance between all pairs. We have incorporated additional distance- and time-based rules for connecting detections to tracks that make the algorithm more robust to missing or false detections by the detection model or from individuals entering or leaving the camera’s field of view during the observation (see Step 3, in the Supplement).

The initial process of linking positions is automated but can result in multiple partial trajectories for a single individual in the case of environmental occlusions and other detection issues. For these instances, we provide a **graphical user interface (GUI)**for easy track validation and error correction. This allows the user to obtain, with limited manual effort, human-verified continuous trajectories of all individuals in each video within the pixel-based coordinate system of the video frame, which can then be transformed to a geographic coordinate system.

### Step 4. Landscape Reconstruction and Geographic Coordinate Transformation

Transforming trajectories into precise geographic coordinates allows us to disentangle the movement of the tracked animals over variable terrain and the movement of the drone (Fig. 3), and also allows the user to analyze the resulting movement data in standard units and in relation to external georeferenced data sources, such as satellite imagery. To achieve an accurate transformation, we must quantify both the topography of the landscape over which the animals are moving and the location of the camera relative to that landscape over the course of the observation (see Supplemental Fig. S1 for an illustration). We obtain this information by first estimating background movement in the video and then using this information to automatically select a subset of frames (typically numbering a few hundred, depending on the extent of camera motion) that capture different but overlapping views of the observation area. We then feed these **“anchor frames”**into **Structure-from-Motion (SfM)**software (Supplement section 4.1). This builds a detailed 3D model of the landscape surrounding each observation and also calculates the location of the camera when each anchor frame was captured. To accurately locate and scale the model in geographic space, we georeference it using either **ground control points** collected in the field or information from the drone’s onboard GPS sensor (Supplement section 4.2). We track visual features in each frame to estimate local camera movement between anchor frames, allowing us to calculate the camera position for every frame in the video (Supplement section 4.3). We then transform animal positions from the pixel coordinates of the frames into the geographic coordinates of the 3D model by projecting rays from the estimated camera location to the surface of the landscape model (Supplement section 4.4). In our examples, this method yields movement trajectories with sub-meter mean and median error (Fig. 3E); we provide a GUI that allows the user to efficiently assess the accuracy of their own track locations (Supplement section 4.5).

**Figure 3.**
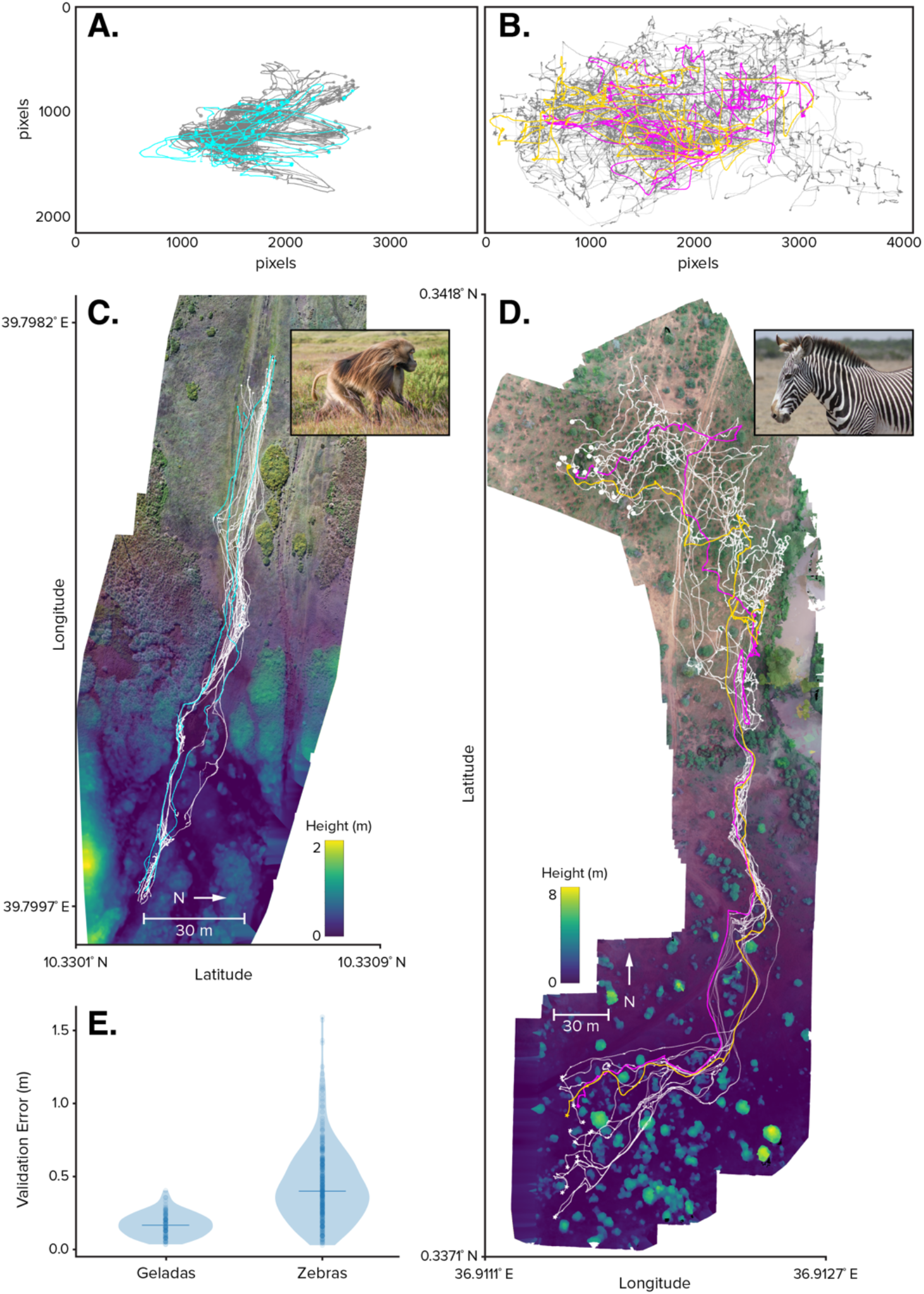
Trajectories extracted from drone videos. Trajectories of A) a gelada monkey group over a period of 5 minutes and B) a Grevy’s zebra herd over a period of 50 minutes, 24 seconds plotted in video coordinates. Drone and animal movement are entangled and the position of the animals on the earth is unknown. In C) and D), the same trajectories are plotted in geographic coordinates and embedded in a reconstructed landscape. The landscape reconstruction includes visual information as shown in the upper parts of map C) and D) as well as topographic information as shown at the bottom. E) The trajectories have submeter human-validated error between observed locations in video frames and embedded map locations (0.17m median, interquartile range 0.11-0.23m from 100 validated gelada locations; 0.40m median, interquartile range 0.26m-0.58m from 350 validated zebra locations). In A and C, light blue tracks denote adult male geladas. In B and D, two tracks are highlighted in yellow and pink for easier comparison. In E, the horizontal line shows the median.

### Step 5. Body-part Keypoint Detection

A major advantage of image-based techniques is that each image contains much more behaviorally-relevant information than does position alone. If the video resolution is sufficient, the user can, for example, use existing tools for markerless pose estimation (e.g. SLEAP (Pereira et al., 2020), DeepLabCut (Mathis et al., 2018; Nath et al., 2019), DeepPoseKit (Graving et al., 2019)) to extract positions of user-defined body parts on each detected animal, generating time-varying postural information for tracked individuals (Step 5 in the Supplement; Fig. 5B). Such postural data are particularly amenable to automated behavioral annotation, and other downstream tasks, because the user can pair the (relatively) low dimensional keypoint information with human verified annotations from the raw video.

**Figure 5.**
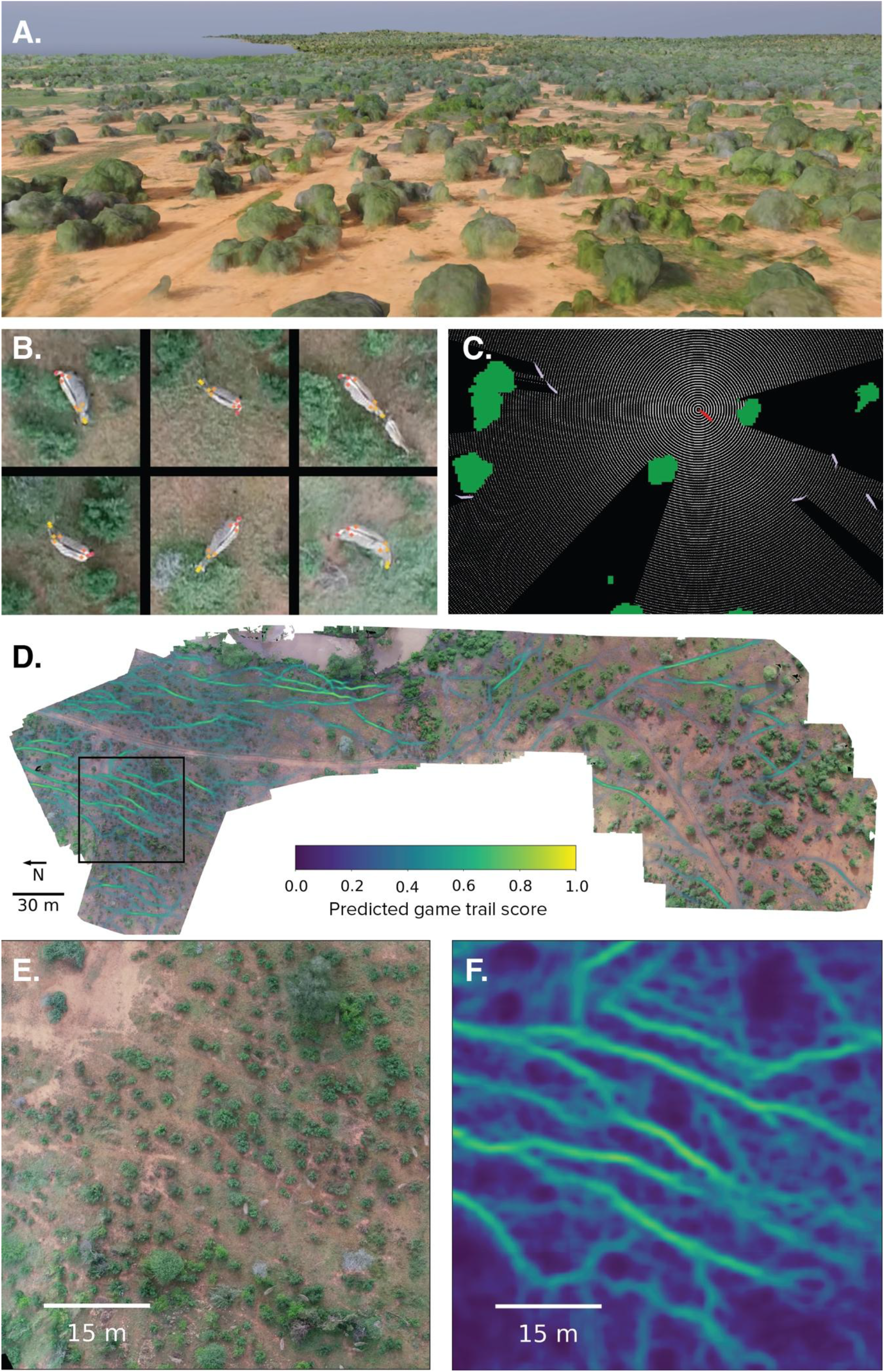
Additional landscape and behavioral data generated beyond animal location. A) An SfM-generated 3D triangular mesh landscape model. B) Examples of keypoints detected on zebras. Nine user-defined keypoints (snout, head, base of the neck, left and right shoulder, left and right hindquarters, tail base and tip of the tail) are tracked using DeepPoseKit (Graving et al. 2019). C) Combining landscape data and animal body and head location allows the visualization of animals’ possible visual fields in their landscapes. Here, the red object is the focal zebra, the white rays show its maximum possible visual field, and the green and white polygons represent bushes and conspecifics, respectively. D) A SfM-generated 2D orthomosaic image showing predicted animal trail presence score. Predictions are generated by a CNN trained on separate orthomosaics that were annotated by three independent human annotators. See worked examples for more detail. The black box indicates the area shown in E and F, which provide detailed views of the orthomosaic and the animal trail predictions, respectively. The color bar applies to both D and F.

### Step 6. Landscape Quantification

The SfM software used in Step 4 generates multiple forms of landscape data that provide valuable environmental context for the behavior of the observed individuals, including 2D rasters encoding elevation data, an **orthomosaic image**, and 3D **point clouds** and **triangular meshes** (Fig. 5A, E). These outputs can be used directly or further processed to extract information regarding local landscape features and topography, classify habitat types, or even estimate the visual fields of tracked animals. In the ungulate worked example we apply pixel-wise classification algorithms, known as **semantic segmentation**, to orthomosaic images to automatically detect possible animal trails, which have been shown to influence the movement of certain savannah dwelling animals (*Cusack et al., 2015; Strandburg-Peshkin et al., 2017*) and are hypothesized (*Kashetsky et al., 2021*) to influence animal movement across a range of contexts (Fig. 5D, F). Further possibilities are explored in the section *Analysis, Extensions, & Applications*.

## Limitations and Considerations

Here we discuss the logistics of using drones to capture behavioral data, the suitability of different research questions to this approach, and the coding skills necessary to implement this method. For required computing resources, please see the Supplement.

### Ethical, logistical and legal considerations

It is important to consider the potential impact of the drone on the focal animals when planning future research. Behavioral and physiological responses to drone flights can negatively impact wildlife (Ditmer et al., 2015; Weimerskirch et al., 2018), and may lead to biased or misleading results in behavioral studies (Duporge et al., 2021). Most species that respond to drones seem to be primarily affected by the sound of the drone (Duporge et al., 2021; McEvoy et al., 2016). Pilots may be able to reduce disturbance by choosing quieter drone models, using low-noise propellers, launching the drone far away from target animals, approaching from downwind, and flying at higher altitudes. While still an emerging field of study, Duporge et al. (2021) and others (Bevan et al., 2018; Christiansen et al., 2016; Mulero-Pázmány et al., 2017) offer guidelines for flying drones near a variety of species. It is typically prudent to perform preliminary flights prior to data collection in order to assess the animals’ response to the drone and establish appropriate protocols.

When choosing a study environment, factors such as extreme temperatures, dust, wind, precipitation and fog can reduce visibility, equipment longevity and flight performance. Furthermore, landscapes (such as open water and snow fields) without abundant and persistent visible landmarks are not suitable for SfM methods, and may preclude transformation of animal trajectories into geographic coordinates (Step 4). SfM height mapping accuracy is also sensitive to vegetation structure and motion (such as tall grass in the wind), which may introduce errors in those parts of the landscape relative to static regions (D’Urban Jackson et al., 2020). Additionally, the real-world geospatial accuracy of the final data is ultimately determined by the quality of the 3D landscape models. For sub-meter accuracy one must either be able to place ground control points in the landscape of interest or have access to existing georeferenced landscape models or imagery.

Finally, researchers must be aware of and abide by legal regulations regarding drone operations in any prospective study area both in terms of formal permissions and licenses required to deploy drones as well as limitations on in-flight maneuvers. Particular rules vary by location and are continually evolving but can include limits on flight altitudes, distances, or locations (i.e. airports, national parks, or certain government areas) and requirements for maintaining visual contact with drones and keeping safe distances from people and structures. Beyond legal requirements, researchers should also ensure that projects do not negatively impact local communities. For a thorough discussion of ethical drone use, see Duffy et al. (2018).

### Research question suitability

In determining whether image-based data collection is appropriate for a research question, researchers should consider the required data resolution and spatial and temporal scales of all targeted study behaviors. Animal groups that are spread over large areas may be impossible to fully capture at an appropriate resolution, especially if automated tracking or individual posture data are required. Deploying multiple drones simultaneously can increase spatial coverage, but complicates flight protocols and data processing. Additionally, spatially-expansive behaviors like long-distance hunts may draw the drone beyond the legal operation distance from stationary pilots (Creel & Creel, 1995). Nocturnal or crepuscular animals would require imaging via thermal or high-sensitivity cameras, which imposes further constraints to the spatial, and temporal, resolution of video that can be obtained. Nocturnal operations also often require additional operation permissions and specialized pilot training.

The timescale over which behaviors occur is also important. Drone battery efficiency has rapidly improved over the years, but flight times are still relatively short (currently 30-45 minutes for widely-used models). While observation time can be extended with sequentially overlapping flights by pairs of drones, theoretically allowing limitless observation times, the practical duration of observations are still limited by battery supply, pilot fatigue, and the movement of the focal animals away from the operational site. Recording rare or unpredictable behaviors, such as predation events, requires deploying drones in the right place at the right time. Incorporating external observational or biologging data could help predict such behaviors and inform the location and timing of deployments. Relatedly, deploying a small number of geolocators on individuals in the target population could allow researchers to more reliably locate rare species or repeatedly target focal individuals and associated conspecifics for drone-based observation.

### Programming proficiency

All code provided in the worked examples is written in the Python programming language. We expect limited additional programming will be required to apply this code to new datasets that are similar in scope to the examples. However, we expect many researchers will want to adapt our base code to better suit their needs, which will require some degree of programming skill, depending on the functionality required. Furthermore, since this method generates large volumes of high-resolution and high-dimensional data, certain programming capabilities will prove essential for effective visualization and analysis (see below for possible analysis directions).

## Analysis, Extensions, & Applications

With recent advances in imaging technology and video analysis, drone-based behavioral observation is poised to become a widely-used approach in the study of wildlife ecology. Here we expand on the potential of this approach by identifying avenues of research that are particularly well-suited to this approach, outlining possible data analysis approaches, and discussing future extensions of drone-based observation.

### Potential research questions

Drone-based observation and our analytical pipeline open the door to several types of questions that can be particularly difficult to explore using existing methods, such as direct observation, camera trapping, or biologging:

1. Our approach allows for precise quantification of the positions of all observed individuals and environmental features, thereby lending itself to exploring *spatially-explicit social behavioral processes*. For example, geladas, plains zebras, and many other animals form multi-level societies, which are reflected in individual association patterns and spatial configurations of groups (Couzin, 2006; Couzin & Laidre, 2009; Grueter et al., 2020; Papageorgiou et al., 2019). These in turn likely affect information flow and collective decision-making, which can potentially be explored by tracking behavioral change and movement patterns.
2. The ability to collect high-resolution behavioral data on all observation subjects simultaneously is conducive to exploring questions relating to *dynamic collective processes*. For example, prey species exhibit vigilance behavior that mediates intragroup information transfer and collective detection of predators (Lima & Zollner, 1996; Lin et al., 2020). Individuals may adjust their vigilance behavior in response to the behavior of group mates, as well as various other social and environmental factors (Beauchamp, 2015; Chen et al., 2021; Pays et al., 2007), but it is challenging to track the behavioral states of multiple individuals simultaneously using non-image-based observational methods.
3. The ability to observe animals and their surroundings concurrently allows for more detailed understanding of *animal-environment interactions*. For example, biologging studies have found that roadways strongly affect wildlife movement, leading to the construction of over- or underpasses to facilitate safe crossing of roadways and promote landscape connectivity (Shepard et al., 2008; Simpson et al., 2016). Aerial cameras could potentially be used to monitor these structures to determine how the presence, speed, and visual characteristics of vehicles affects animals’ use of these structures.

To the extent that these examples can be addressed using other methodologies, they will require multiple sensors, fusion of diverse data streams, and/or considerable effort by human observers. In contrast, drone-based observation allows for all the relevant data to be collected simultaneously using a single sensor.

### Data analysis

Our method provides a rich set of biologically-relevant features, but these data must be analyzed to address the research questions of interest. There exists a rich body of literature for exploring, visualizing, and analyzing animal trajectory data (Patterson et al., 2017; Seidel et al., 2018). The sub-second and sub-meter animal positions and detailed landscape data produced by this method allow researchers to use step-selection-type approaches to evaluate movement decisions across spatial and temporal scales, from actual individual steps to movement decisions at larger-scales (Fieberg et al., 2021). Tracking body keypoint locations (Step 5) can directly provide information about behaviorally-relevant body parts, such as head direction and body orientation. Depending on the position of the animal(s) within each frame, correctly interpreting animals’ relative keypoint patterns may require taking lens geometry into account (see Supplemental Fig. S3 for details). When possible to estimate, this head and body information can be combined with the 3D landscape models (Step 4) to reconstruct estimates of each animal’s visual field (Aben et al., 2018).

### Future extensions

During detection (Step 2), if given sufficient image resolution, it may be possible to visually identify individual animals within and across observations. Individual recognition opens the door to studies of individual behavioral variation across time and contexts, and the role of individual behavior in driving processes at the population and community levels (Costa-Pereira et al., 2022). Beyond laboratory settings (Walter & Couzin, 2021) individual recognition in wild populations is increasingly feasible (Norman et al., 2017; Tuia et al., 2022). Alternatively, individuals could be identified using ground-level photographs or direct observation, and then these identities could be manually linked to individuals in the aerial recordings.

A further advance would be to use temporal keypoint data, or body posture trajectories (Fig. 1 - Step 5), to define finescale behavioral labels. While similar to the problem of accelerometer-based behavioral classification, a vision-based approach provides the added benefit of ground-truth videos for validation (Brown et al., 2013; Wang et al., 2015). Both supervised (Bohnslav et al., 2021) and unsupervised/self-supervised (Berman et al., 2014) approaches, or a combination of both (J. J. Sun et al., 2021), could be applied to achieve automated behavioral annotations describing the behavioral states of individuals and groups.

Beyond data processing, advancements with drones’ on board automated visual tracking (Islam et al., 2019) and the ability to automatically coordinate flight among multiple drones (Zhou et al., 2022) could help to streamline complex operations, reduce the risk of human error, and also facilitate further observation techniques such as multiview 3D posture tracking (Tallamraju et al., 2019). Drones could thus be deployed to autonomously find and record individuals of a target species. While these are exciting future steps, current regulations in many countries would prevent the use of these methods without special permissions or certifications. Thus, regulatory rather than technological restrictions may be the most substantial barriers to large-scale automated observation.

### Synergizing with other remote-sensing methods

There is great potential in combining this approach with existing remote sensing methods. In the context of biologging, drones could be deployed at key times or locations of interest to provide high-resolution behavioral snapshots of tagged individuals along with their social and environmental context. Image-based data could also be used to guide biologging study design choices. For example, one could downsample the high resolution image-based data to determine the optimal sampling frequency for bio-logging studies, using the videos to verify that the resulting data capture the target behaviors. Recording the behavior of instrumented animals could also provide sensor ground truth data and aid in the development of more energy efficient, behaviorally activated sensors (Korpela et al., 2020; Yu et al., 2021).

The geolocated animal movement and environmental data generated by drone-based methods can be combined with additional multi-modal remote sensing data to explore the interplay between animals and important environmental features. For example, higher accuracy canopy height and landscape models, such as those generated by emerging workflows of SfM, LiDAR, or their fusion (D’Urban Jackson et al., 2020), combined with animal behavioral data generated from our method, could enable studies of herbivore impacts on vegetation via consumption or trampling effects, or allow higher precision visual field calculations in difficult-to-model environments like tall grass. Additionally, SfM models, such as those generated in our method, can be combined with various other remote sensing data to enable calculation of microclimate variability (Duffy et al., 2021; Maclean, 2019) water flow and saturation (Koci et al., 2020), or resource quality and accessibility (Jennewein et al., 2021) which would also be informative for analyzing recorded animal behavior.

## Conclusion

We present a method that allows researchers to study animal behavior in its natural social and environmental context in a non-invasive and scalable way. Our approach is independent of specific species and can be deployed across a range of study systems making this a powerful and versatile tool for many researchers across fields. Importantly, as researchers work to understand the relationship between animals and their landscapes in a changing world, this method, which simultaneously records both at high resolution, is poised to be an important new way of observing the natural world.

## Authors contributions

Benjamin Koger, Blair Costelloe and Iain Couzin developed the idea and goals of the method. Benjamin Koger, Blair Costelloe and Jeffrey Kerby coordinated and conducted data collection. Benjamin Koger, Blair Costelloe, Adwait Desphande, Jacob Graving and Jeffrey Kerby curated the data. Benjamin Koger designed the analytical pipeline; wrote the code in the worked examples and performed all aspects of model development, including training and validation. Jacob Graving advised on the design, organization, and development of earlier versions of the code. Blair Costelloe and Iain Couzin provided supervision and secured financial support. Blair Costelloe, Iain Couzin and Jeffrey Kerby administered and contributed material resources to the project. Benjamin Koger and Blair Costelloe wrote the initial manuscript draft and prepared the figures. All authors edited and revised the manuscript and gave final approval for publication.

## Acknowledgements

Figures 1 and 4 were created by Mike Costelloe. We thank Mike Costelloe, Sven Lauke and Felicitas Oehler for assistance with collecting and annotating the ungulate data. We thank Guassa Gelada Research Project field managers Odin Bernardo and Nate Redon for collecting the gelada drone data, and project directors, camp staff members, and the local community for their support. We thank Anne Scharf for sharing code used to plan DJI mapping flights. We thank Jean Luo and David Basili for contributions to prior versions of the animal trail detection and track correction GUI. B.R.C. received support from the European Union’s Horizon 2020 research and innovation program under the Marie Sklodowska-Curie grant agreement No. 748549. B.R.C. acknowledges support from the University of Konstanz’s Investment Grant program. J.T.K acknowledges support from the Neukom Institute for Computational Science at Dartmouth College and the European Union’s Horizon 2020 research and innovation program under the Marie Sklodowska-Curie grant agreement No. 754513 and the Aarhus University Research Foundation. B.K., I.D.C, J.M.G., B.R.C., and A.D. acknowledge support from the Deutsche Forschungsgemeinschaft (DFG, German Research Foundation) under Germany’s Excellence Strategy – ‘Centre for the Advanced Study of Collective Behaviour’ EXC 2117-422037984. I.D.C gratefully acknowledges support from the Office of Naval Research (ONR) grant N00014-19-1-2556 and the European Union’s Horizon 2020 research and innovation program under the Marie Sklodowska-Curie grant agreement No. 860949. A.D. acknowledges the support by the Swiss National Science Foundation (Early PostDoc Fellowship: P2NEP3_200190). J.M.G. and B.R.C. acknowledge support from NVIDIA Corporation’s Academic Hardware Grant Program.

## Permissions and Ethics Statement

We imported and operated drones in Kenya with permission from the Kenya Civil Aviation Authority (import permits: KCAA/RPA/PERMIT-2017-0006, KCAA/RPA/PERMIT-2017-0007, KCAA/RPA/PERMIT-2017-0008, KCAA/ASSR/RPA/PERMIT-0016, KCAA/ASSR/RPA/PERMIT-0017, KCAA/ASSR/RPA/PERMIT-0018; authorization numbers: KCAA/OPS/2117/4 Vol. 2 (80), KCAA/OPS/2117/4 Vol. 2 (81), KCAA/OPS/2117/5 (86), KCAA/OPS/2117/5 (87); operator certificate numbers: RPA/TP/0005, RPA/TP/0000-0009). We conducted research in Kenya with permission from the Kenyan National Commission for Science, Technology and Innovation (research permits: NACOSTI/P/17/59088/15489, NACOSTI/P/59088/21567) and in affiliation with the Kenya Wildlife Service. All research activities pertaining to the ungulate data, including drone operation, were performed with the knowledge and support of management and security staff at our field sites, Ol Pejeta Conservancy and Mpala Research Centre. Data collection protocols for the ungulate data were reviewed and approved by Ethikrat, the independent ethics council of the Max Planck Society. All research activities pertaining to gelada monkeys, including drone operation, were undertaken with the knowledge and approval of the Guassa Community Conservation Area leadership and under the approval of a memorandum of understanding between the Ethiopian Wildlife Conservation Authority and the Guassa Gelada Research Project.

## Data availability statement

Code for the worked examples is available on GitHub at https://github.com/benkoger/overhead-video-worked-examples. All data for the worked examples are archived on EDMOND at https://edmond.mpdl.mpg.de/privateurl.xhtml?token=9c1a978a-21f3-4843-bebe-40c296bffc73. DOIs and stable links to these resources will be generated upon final acceptance of the manuscript.

