## Supplemental Text for "Quantifying the movement, behavior, and environmental context of group-living animals using drones and computer vision"

### Details of the method

*Step 1. Video Recording*

Our method assumes that terrestrial animals are recorded from overhead with the camera pointing directly down. This nadir viewpoint reduces occlusions from vegetation and other animals, results in a relatively uniform appearance of all individuals regardless of their body orientation, and simplifies the task of calculating camera movement. Good overall visibility of the animals is fundamental to the detection and tracking method: a good criterion for this is that objects that are readily detectable by humans when viewed in full resolution are typically also detectable by a well-trained model. Continuous tracking will be easiest in habitats with sparse canopy cover, but brief occlusions from passing under vegetation are generally not difficult to correct using our provided Graphical User Interface (see Step 3, below). Greater visual contrast between the animals and the ground allows for easier detection by the model and requires less human annotation effort (see Step 2, below). Cryptic or very densely clustered animals such that the boundary between individuals is not visually distinguishable may be challenging to annotate and are likely to not be reliably detected by the trained model without special effort.

An important factor affecting detectability is the size of the animals in pixels in the video frames. While there are no hard thresholds for minimum object size, as a guide, spatial resolution should be high enough to allow for easy human visual detection of the animals and any relevant body parts for posture tracking (see Table S1 for information on the size in pixels of animals in the worked examples and corresponding model performance). Required resolution may change seasonally with changing coloration of animals or landscapes. Most cameras have a range of high- and ultra-high-definition settings, typically (at the time of writing) being at least 4K, or 3840x2160 pixels; higher definition settings will result in more pixels per object, but will also increase file size of the videos and processing time for the detection model (see Step 2, below). In addition, there exists an inevitable tradeoff between spatial resolution and spatial coverage: as spatial resolution increases, either by moving the camera closer to the ground or zooming in, the spatial coverage of the video frame decreases. This may limit the number of individuals that can be filmed simultaneously and make it more difficult to keep all individuals in the frame, especially if animals are spread out or moving quickly, which may also prevent continuous tracking and maintenance of individual identities throughout the observation. Furthermore, when recording with drones, zooming in may only be possible by flying at lower altitudes, which may negatively affect the animals (see Limitations and Considerations in the main text). Note that digital zoom (as opposed to optical zoom) effectively captures information from a smaller region of the image sensor, potentially resulting in a reduced video sharpness and quality. It is important for the user to carefully consider the necessary image resolution of their subjects for the specific research task so that this can be achieved without unnecessarily increasing processing time, restricting spatial coverage, or disturbing study subjects.

Video should be recorded at a high enough frame rate to capture the behaviors of interest, and to ensure high quality tracking (see Step 3, below) which depends on the distance between an animal's location in consecutive frames being less than the distance to its neighbors. As a lower bound on framerate, behavior must be recorded at at least twice the frequency of the behavior of interest (Shannon, 1949). In our worked examples thirty frames per second (fps) were sufficient. Unnecessarily high frame rates should be avoided as the resulting videos will require more data storage capacity and drastically increase processing time.

*Step 2. Detection and Localization*

Convolutional neural networks (CNNs) are a standard approach for detecting and localizing objects in complex images and we use them here to detect and localize animals in each video frame. To train a CNN, one must create three sets of **annotated** images (see Table S2 for definitions of bolded terms) in which all objects of interest have been labeled, for example by drawing bounding boxes around each animal. The training set is the largest set and provides the examples from which the model learns which objects to detect and what background imagery to ignore. The validation set is also used during training but to evaluate the model as it learns and to determine whether changes to the model improve or impair performance. The test set is used only after all training has finished to measure the performance of the final model on images it has never seen before. Annotation can be tedious and the content of the image sets will strongly affect model performance; therefore, it is important to carefully consider the annotation strategy in order to achieve high information value in the image sets while minimizing human labor (see 2.1 Image annotation, below).

In selecting a detection model, the user can choose from a range of options that will give good results with various speed and accuracy trade offs (see Huang et al. (2017) for a detailed comparison of various standard models). For simple use cases, where the animals are easy for a human to see in the videos, many common models can be readily configured for the researchers' data (see 2.2 Model training, below). Researchers with more complex detection requirements may have to move beyond standard models and training regimes. As a starting point, we describe the approach we use in our worked examples, which should yield suitable results for simple use cases.

*2.1 Image annotation*

A common question regarding image annotation is how many examples are needed to train a given model. In reality, the answer is a function of both the complexity of the videos and the desired quality of the ultimate model. In cases where animals are easy to see, a model trained on only a few hundred annotations (note that there could be multiple annotations in one image) will detect many individuals. For truly robust results in a range of habitat conditions annotating a few thousand individuals is a better starting target, although in particularly challenging cases the number may be higher (see Table S1 for number of annotations used in the worked examples). More important than the raw number of annotations, however, is the overall information content of the training set and how well those annotations reflect the intended natural conditions of all observations. Training sets with many very similar annotated examples are less valuable than a training set with more variable examples. In particular, if the annotated examples used in the training set reflect just a subset of the observation conditions the result could be a model that does not generalize well to the full set of observations included in the study (see 2.2 Model training, below, for tips to mitigate this). The researcher should choose images that show the target animals in the full range of different landscapes and poses that appear in the complete set of observations.

Importantly, the images in the training, validation, and test set should be independent examples of the overall expected observation conditions. This is driven by the challenge that, with too much training, the model will overfit to the training data, meaning the model relies on attributes that are specific to the training images but that do not generalize across examples in the whole dataset. As long as the validation and test sets provide uncorrelated samples of the expected distribution of conditions, overfitting can be detected and prevented. Conversely, if the subset of images used to build the validation set is similar to those used to build the training set, but does not encompass the complete data set, then overfitting is hard to detect.

Building diverse and independent annotated image sets from video recordings presents a challenge because video frames are temporally autocorrelated: nearby frames can be nearly identical, particularly when filming slow-moving animals at high frame rates. Similarly, there is likely strong visual correlation between annotated objects within a single video frame, as they will be set against similar backgrounds and subject to similar lighting conditions. This is of particular concern when there are many objects of interest in each frame – for example when working with recordings of large groups of animals – since annotating such frames is time-intensive and the added informational value of each additional annotation in the frame quickly diminishes.

Users can avoid autocorrelation across frames by randomly selecting frames from the dataset for annotation, rather than annotating sequences of consecutive frames. To maximize independence of the training, validation, and test sets, entire observations should be randomly assigned to one of the three sets before randomly selecting frames from each observation for annotation. If there are insufficient observations for this approach, each observation can be split into segments, with time buffers between each segment. These segments can then be randomly assigned to one of the three sets before frame selection. Non-overlapping sets are most important in cases where the researcher wishes to investigate possible model performance on completely unseen observations (e.g. data to be collected in future field seasons). If all observations are already in hand, it is sufficient to randomly sample frames from across the entire dataset and then randomly assign each frame to one of the three image sets. Generally, the majority of images are put in the training set with fewer in the validation and test sets. Standard ratios are 80% of images in the training set (Goodfellow et al., 2016), but it depends on the number of images annotated and the variability of the visual conditions. To reduce the visual correlation that can result from annotating many objects in the same frame, users can annotate cropped portions of frames instead of entire frames. Note that it is important to avoid cropping too tightly around objects of interest in order to include substantial background imagery so that the model can also learn what should be ignored (e.g. large rocks, or landscape driven shadows). The minimum possible crop size is determined by the input requirements of the model. In the gelada worked example we use 1024x1024 pixel crops while we annotate full frames with the ungulates.

Once the images for annotation have been chosen and allocated to the training, validation and test sets, the user must choose an annotation tool to actually label the images. While the offering of possible annotation tools is continuously evolving and too dynamic to meaningfully review here, there are a host of options that vary in complexity, cost, and functionality. Aspects to consider are if the tool is free to use and open source (such as CVAT (CVAT.ai Corporation, 2018/2022), AIDE (Kellenberger et al., 2020), the VGG Image Annotator (Dutta & Zisserman, 2019), or LabelImg (Tzutalin, 2015)) or is a commercial product (such as Labelbox (2022)) which may require payment but may also provide more customer support. We use Labelbox (2022) with a free educational license. Most image annotation tools, including all mentioned above, allow 'bounding box' annotation, which is what we use. Note that many tools also allow the user to annotate sequences of consecutive or nearby video frames. This speeds up annotation since less searching is required to locate all individuals in complex images and bounding boxes drawn in the first frame of a sequence need only slight adjustments in subsequent frames. However, there is a tradeoff between increasing annotation speed in this way and maintaining high information content in annotations as frames close together in time will have high information correlation.

Another technique to improve the efficiency of the annotation process is **model-assisted labeling**, an iterative process where image annotation and model training are conducted in parallel. The user initially labels only a subset of the images in the training set and uses these to train an initial version of the model. This initial model is then used to pre-annotate the remaining images. Further annotation then involves confirming, or fixing, the model derived pre-annotations for each frame while also adding annotations for any animals that the model missed. This can significantly speed up annotation (Pereira et al., 2019) since the model will quickly learn to detect the majority of easier cases but tends to struggle with a minority of hard cases that the human can correct. Additionally, since the trained model generates confidence scores for each annotation, the user can see which objects are particularly easy or hard for the model to detect confidently. Here, it is useful to incorporate an **active learning** (Kellenberger et al., 2020; Tuia et al., 2011) approach where, after the first round of model training, annotation effort is concentrated on annotating images with low confidence scores, reducing redundant manual labor and allowing the model to learn more efficiently.

#### *2.2 Model training*

To successfully train a deep learning-based object detection model, researchers must choose an appropriate software framework, a specific model to train within that framework, and, finally, the specific protocols for training that model. These choices will be driven by the researcher's experience with deep learning tools and the complexity of the recorded videos and so two researchers may make different choices to meet their needs.

The deep learning framework is the software tool or programming library that interfaces between the user and the computer to define, train, and use a deep learning-based object detection model. There are commercial entities such as Datature (https://datature.io) and LoopBio (http://loopbio.com) that offer code-free solutions, but also many robust open-source coding-based frameworks such as PyTorch (Paszke et al., 2019), Keras (Chollet & others, 2015), and Tensorflow (Martín Abadi et al., 2015).

Importantly, these open-source application programming interfaces (APIs) allow the user to interact with model components at different levels of abstraction so that common models and training regimes can be deployed quickly with minimal coding, while users with the necessary expertise can customize when necessary. Importantly, major frameworks have prebuilt implementations of common object detection models (such as Faster-RCNN (Ren et al., 2015) and Retinanet (Lin et al., 2017)). While standard modifications are necessary to apply these models to new datasets, as explained below, the researcher can rely on the pre-built models for core implementation details. We demonstrate using the PyTorch framework with Detectron2 (Paszke et al., 2019; Wu et al., 2019) in the worked examples.

CNN-based object detection models vary in performance in terms of speed, accuracy, and memory requirements. While new models are always being developed, most researchers will choose one that is already implemented in their chosen framework and that matches their performance requirements. This both decreases coding time and allows training with smaller training sets, as described below. Object detection models are built around a backbone which translates raw image inputs, where each pixel encodes red-green-blue color information (if typical video cameras are employed), into an abstract output, where each spatial point corresponds to a high-dimensional vector (a few hundred or thousand dimensions) that encodes features like texture or object shape. The backbone is built from stacks of spatial convolutional filters, non-linear transforms, and spatial pooling layers that compress local spatial information together. More layers allow the model to learn more complex relationships that span more of the image, but require more memory and incur slower processing speeds. The output of the backbone is then used by the model to localize and classify objects in the image. Some models, like SSD (W. Liu et al., 2016) and YOLO (Redmon & Farhadi, 2018) perform this step more quickly but with lower performance while others, like Faster-RCNN (Ren et al., 2015) are more accurate but are slower. Deep learning frameworks with pre-built models include information about the performance and processing speed of each available model. While the actual values will be specific to the data they were trained on (for accuracy measures) and the hardware that was used (for speed measures), they serve as a useful relative indication of expected performance and speed/accuracy tradeoffs. In the worked examples we use Faster-RCNN (Ren et al., 2015) with a Resnet 50 Feature Pyramid Network backbone (Lin et al., 2017).

General purpose deep CNNs used for object detection in complex scenes, such as ours, can have millions of parameters that must be learned. Training such a model from scratch requires many thousands of training images and large amounts of computational time (He et al., 2019). Instead, researchers commonly fine-tune the weights of a model that has been pre-trained on a large public dataset containing images and annotated objects across a range of common conditions (Li et al., 2019). This pretraining teaches the model the basic parameters of the visual world and generates initialization values for the model weights that are then trained on the user's smaller dataset. Frameworks for object detection provide pre-training weights for their included models. In our case, the models we used were pre-trained on the Common Object in Context (COCO) image dataset (Lin et al., 2014).

After the researcher has chosen a model, they must ensure it is configured to work well with their specific data. One necessary modification configures the model to choose among the researcher's possible output classes for each detected object. In the model, this selection happens at the last layer and is done with multinomial logistic regression. The input to the regression is the abstract high dimensional representation of the contents of the bounding box that is a fixed size regardless of the number of classes being predicted and the output is an N dimensional vector where N is the number of possible classes in the researchers' dataset. A second task is to ensure the model training pipeline is configured to efficiently read in the training images and annotations. Large images take a long time to move between a computer's memory and the processing cores on the CPU and/or the

GPU. Deep-learning frameworks provide efficient pipelines for speeding up this process but expect either the images and annotations to be in specific data formats or for the user to write their own data loaders that reflect their data structure. The user must either modify their training data to fit the expected format or modify the training code to match their data. The specific formats depend on the framework being used, but inefficient data loading can be a bottleneck when training and using deep learning models (see worked examples for examples with Pytorch and Detectron2). During training, if the computer shows a low GPU's utilization measure this can be a sign that data loading is a bottleneck.

Another important part of training is **image augmentation**, a common component of the input pipeline where images and their annotations in the training set are modified each time they are passed to the model (Shorten & Khoshgoftaar, 2019). These modifications, for example mirroring or blurring the image and corresponding annotations, reflect the expected image variation of the full dataset. As an example, adjusting the brightness and contrast of a training image can simulate a particularly dark or cloudy day. In practice, each time an image is loaded one or many augmentations are applied to it with the degree of augmentation defined by a random distribution. Image augmentation increases the effective size of the training set without the need for additional human annotation, allowing for better model generalization (Shorten & Khoshgoftaar, 2019). In our worked examples we include horizontal and vertical mirroring and brightness and contrast modification. Most training pipelines have some common image augmentation built in and there are many packages such as imgaug (Jung et al., 2020) and albumentations (A. Buslaev & Kalinin, 2018) available for adding diverse augmentations.

Finally, once the training pipeline is in place, the researcher must evaluate how long to train the model on the annotated data. The model's performance on the validation set is used to determine training duration. After a specified number of training iterations, often after all images have been shown to the model once (called an epoch), the validation set is fed to the model to evaluate the model's performance on images it has not learned from. The model should be trained as long as the performance on the validation set continues to improve. While performance on the training set may continue to increase after this point, those gains are driven by overfitting (see Table S1 for performance metrics for the trained models used in the worked examples).

When training a model there are further possible hyperparameter choices, for instance about the model's specific loss function, learning rate, and optimization algorithm, that affect the ultimate performance of the model. However, one needs an understanding of the details of deep learning that goes beyond the scope of this paper to decide when specific choices beyond the default make sense. Instead, we suggest that if one does not have that experience, focus should be given to obtaining videos where animals are easy to see so standard implementations will give good results.

After the model is trained, all required video frames from all required observations can be processed; for each video frame the model generates bounding box coordinates, predicted object types, and confidence scores for every object detected. We take the mean of the coordinates associated with each corner of the bounding box as an individual's location in the image.

| **Object class** | **N images (train/test)** | **N annotations (train/test)** | **Bounding box area (pixels) Mean (lower quartile, upper quartile)** | **Precision** | **Recall** |
|---|---|---|---|---|---|
| Ungulate model | | | | | |
| zebra | 1,007/211 | 14,315/3,094 | 3660 (2626, 4218) | 0.99 | 0.98 |
| impala | 378/77 | 4,830/980 | 1604 (1089, 1920) | 0.96 | 0.96 |
| buffalo | 192/47 | 6,192/1,615 | 5650 (3690, 7392) | 0.98 | 0.93 |
| waterbuck | 63/8 | 498/40 | 2888 (2472, 3256) | 1.0 | 1.0 |
| other | 172/35 | 854/169 | 1625 (1363, 1885) | 0.98 | 0.97 |

| | | | | | |
|---|---|---|---|---|---|
| Gelada model | | | | | |
| adult-male gelada | 55/11 | 239/25 | 743 (494, 944) | 0.86 | 0.72 |
| other-gelada | 65/16 | 1821/160 | 393 (240, 496) | 0.89 | 0.81 |
| human-observer | 17/7 | 29/7 | 952 (416, 1443) | 1.0 | 1.0 |

**Table S1: Information about the annotated datasets used in the worked examples and corresponding performance metrics for the trained detection models.** For the performance metrics we used an intersection-over-union (IOU) score of 0.6 and a minimum confidence score threshold of 0.7 for each detection. Precision is the proportion of detected instances that were actually in the target class (true positives / (true positives + false positives)). Recall is the proportion of the actual target class instances that were detected (true positives / (true positives + false negatives)). For the ungulates examples we use the validation set to calculate the performance metrics since we did not build a true test set. While we these numbers may be interesting for researchers thinking about the number of required annotations needed for their project and the size at which to film individuals, we emphasis that there are other factors, such as contrast of animals against background, and the expected variation in environmental conditions, that strongly affect the challenge of the detection task and therefore the size at which individuals should be filmed and the number of required annotations. We recommend looking at our actual annotated images to gauge the challenge associated with each class presented here. Relatedly, here we record the size of the bounding box, not the actual animal. Especially for the ungulates, many of the bounding boxes are larger than the animal it contains. Lastly, we note that we did not train a separate model for each class. As described elsewhere, two models were trained: one for all the ungulate classes and one for all the gelada classes. Training a model on a training set with class imbalances can negatively affect performance on underrepresented classes.

*Step 3. Tracking*

Once the detection model has located the individuals of interest in each frame, these positions must be associated across frames to construct each individual's trajectory throughout the video sequence during which the animal is present. We combine automated tracking with human oversight to efficiently construct verified individual trajectories throughout each recording. We first automatically construct partial trajectory segments from detections, ending segments when there is ambiguity about which detection to connect next. The provided **graphical user interface** (**GUI**; see worked examples) allows the researcher to easily visualize and, if necessary, combine these segments into longer trajectories. The user's specific research questions will determine the level of required human oversight. Research questions that do not require individual identities to be maintained throughout an observation, for instance, may be able to use the uncorrected partial trajectories. Here, we present a simple yet robust approach that uses the distance between detections in consecutive frames to link locations and reconstruct trajectories. We also note, however, that multi-object tracking in video is a quickly developing area of research with many powerful techniques that may increase tracking performance beyond what we demonstrate here (Park et al., 2021).

Most consumer cameras film at at least 30 fps with rates of 60, 120, and even 200 fps becoming possible. Our tracking approach relies on the assumption that the combined camera and animal movement produce only modest changes in individuals' positions from one video frame to the next. In our applications, 30 fps proved fast enough to fulfill this criterion.

If all individuals are detected perfectly throughout a video then distance-based tracking can be done with a simple energy minimization algorithm such as the Hungarian algorithm (Kuhn, 1955). The Hungarian algorithm is designed to solve the situation where there is a collection of new positions that must be uniquely assigned to a matching number of existing trajectories. This algorithm finds the pairing of positions and trajectories that minimizes the total distance between all pairs. In our use cases, however, the requirement of an equal number of trajectories and new positions is often not met because missing or false detections from imperfect model performance lead to a variable number of detected individuals across frames, and, also, because individuals may enter or leave the camera's field of view over the course of the video.

We address these problem cases by supplementing the Hungarian algorithm with a few simple rules. First, we only allow a position to be connected to an individual's trajectory if it is within a distance defined by the lower of either a user-defined global maximum distance or .45 times the distance to the next closest trajectory. Thus, as individuals get closer to each other we get more conservative in assigning new positions to existing trajectories, resulting in fewer cases of trajectories switching between individuals. Because of this upper threshold, we optimize over the natural logarithm of the distance between points, plus one, (ln(1 + distance)) so that false long distance pairs that will ultimately be filtered out by the threshold do not disproportionately influence the global optimization. The second modification we make is that trajectories that are not assigned a new position in a given frame stay active for a few subsequent frames in which they can be assigned new positions if an appropriately located new animal detection appears. If this happens, positions are simply interpolated between the last detected position in the trajectory and the new one. After a certain number of frames with no new position, however, the trajectory segment is considered finished. Relatedly, positions that are not assigned to an existing trajectory are used to create new trajectories. However, if new positions are not added to these new trajectories in subsequent frames, they are considered noise and are deleted. Lastly, if there is only one new point within a track's threshold distance, if that track is longer than nearby tracks, the longest track takes precedence. With this approach, animals that remain visible and well detected by the model generate complete trajectories without human intervention. Animals that are temporarily occluded or only partially detected by the model generate a series of trajectory segments where each segment has a high likelihood of coming from a single individual.

To review and verify trajectories, we provide a GUI that can be used to visualize the trajectories for each individual and efficiently connect segments together. This allows the user to obtain, with limited manual effort, human-verified continuous trajectories of all individuals in each video within the pixel-based coordinate system of the video frame.

*Step 4. Landscape Reconstruction and Geographic Coordinate Transformation*

The trajectories generated in Step 3 only record the movement of individuals with reference to the coordinates of the video frame (Fig. 3). This is a problem because movement in the video space is a function of both drone movement and animal movement. Likewise, distance between animals in video space is also a function of both the drone's altitude, the landscape topography, and the actual distance between individuals. Typically, the researcher is actually interested in the animals' movements and relative positions in the coordinates of the earth (i.e. latitude, longitude, elevation), independent of the above properties of the drone. Therefore, it is essential to project the locations from pixel coordinates into geographic coordinates.

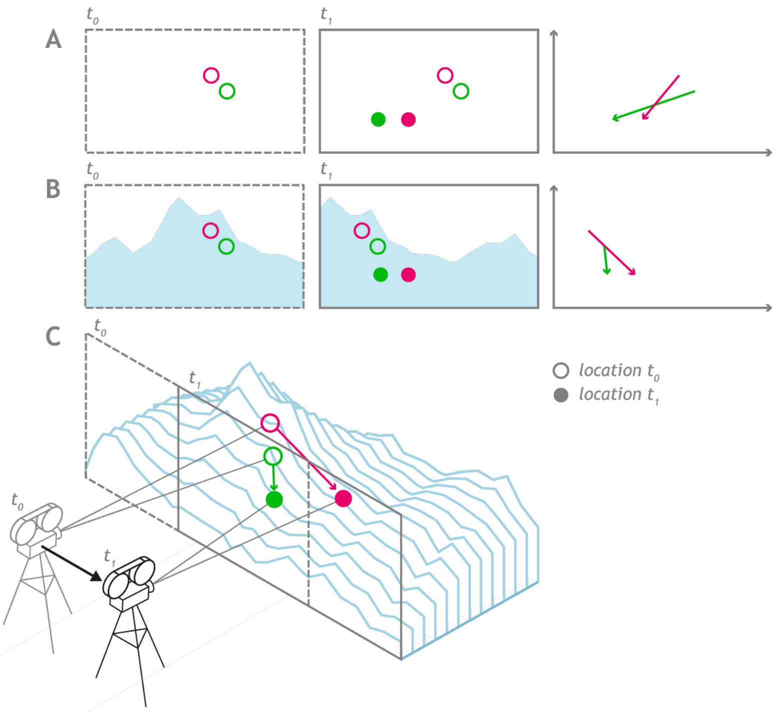

**Figure S1: Both camera position and landscape topography are necessary to generate accurate trajectories from locations extracted from 2D images.** A) At the end of Step 3 in the data extraction pipeline, we have linked detections across frames to generate trajectories for individual animals in the pixel-based coordinate space of the video frames. These trajectories do not distinguish between camera movement and animal movement, and thus changes in animal positions across frames (here from t0 to t1) do not accurately represent animal movements in the real world. B) Adding the environmental context (a mountain imaged from the side) to the images highlights this discrepancy. C) By combining camera location information with the 3D topography of the environment, we can reconstruct accurate movement trajectories in 3D space. Without a model of the landscape structure, the user must assume a flat earth, which introduces error into trajectories. This conceptual figure illustrates the example with a horizontal camera filming a mountainous landscape to more clearly emphasize the role of topography. These principles translate to the nadir camera position of the worked examples.

Since light's path of travel is deterministic, its path from a specific point on the ground, through the camera lens, and onto a given pixel on the camera sensor is fixed for any camera location. As long as the topography of the landscape and the camera's location is known, we can traverse that fixed path in reverse to project the position of a camera's pixel back onto the earth (Fig. 4). Although information about the 3D topography of the landscape and the camera's location in that space is not naturally embedded in a single image, with certain flight protocols (see Step 1, above) and some additional processing we can extract this information from observation videos.

Here we demonstrate how to project movement trajectories from video coordinates to geographic coordinates with a typical sub-meter mean and median error.

*4.1 Using Structure-from-Motion (SfM) to determine landscape topography and camera locations*

We use SfM to create 3D landscape models. This technique matches visual features across a set of overlapping 2D images - in this case, specific video frames - to infer the geometry of the landscape. SfM requires that the same landscape features be visible in many images, thus constraining the geometry of the landscape and simultaneously defining where the camera was relative to this geometry when each image was captured.

SfM is computationally intense and attempting to process all of the tens of thousands of frames from a drone observation can overwhelm many computers. However, due to the high frame rate of video, under normal filming conditions, even a small subset of frames has enough visual feature overlap to generate high-quality 3D landscape models. Although using a subset of frames is necessary to make SfM processing feasible, it also means we only generate camera location information for the frames in this small subset. Therefore, after generating the 3D landscape model, we use the location information for this subset of frames as a basis to estimate camera locations for all other frames in the observation (see 4.3 Estimating camera locations for all frames*,* below). For this reason, we call this subset of frames **anchor frames**.

We choose anchor frames based on the camera's movement as logged by the drone's on-board GPS and accelerometer sensors (see ungulates worked example) or based on estimating local visual movement between video frames (see gelada worked example). To ensure sufficient overlap between images, we add an anchor frame after the drone has moved or rotated beyond a certain distance from the last anchor frame. Beyond image overlap, the final resolution of the landscape model is limited by the imagery's ground sampling distance, or the ground distance between adjacent pixel centers in an image. In addition to adding anchor frames to ensure model quality, error can accumulate when estimating camera location between anchor frames, particularly when the drone moves. Therefore, we also add anchor frames before and after large drone movements and rotations so any errors introduced during movement do not spread to the potentially long periods of hovering before or after.

Once extracted, the anchor frames are passed to a SfM software to calculate the 3D topographic model and camera location information. For the worked examples we use Pix4D Mapper [Educational license, Pix4D SA, Switzerland] but there are many software tools to choose from.

*4.2 Georeferencing the landscape model*

Raw video frames have no associated location information; therefore, the 3D model generated in the previous section has no information about the location of the model on Earth nor about the internal scale of the model. Animal location information can be projected into this landscape model (see next section), but the trajectories will lack geographic coordinates and the scale will be unique to the individual landscape model and observation, precluding incorporation with external datasets or analyses in standard units. Additional georeferencing information must be supplied to the SfM software to add location and scaling information. The resulting geolocation accuracy of the model determines the accuracy of the geospatial data that is ultimately derived from it, including the animal trajectories. Model accuracy can vary in terms of internal accuracy (the accuracy of points in relation to each other in the model) and absolute accuracy (the relationship between points in the model and the coordinates of points externally measured in the landscape). Internal accuracy is important for evaluating things like distance between individuals and an individuals' speed, while absolute accuracy is important for combining animal trajectories with external datasets, such as road maps or satellite imagery.

In the context of drone imaging, the simplest way to georeference the landscape model is via direct georeferencing where location and orientation information for each anchor frame is extracted from the flight logs recorded by the drone's on-board sensors. This method is efficient because no additional information needs to be collected, but it often results in lower quality georeferencing compared to other methods (Kalacska et al., 2020). In particular, generating precise models of long narrow landscapes, a common result when following animals on the move, is known to be a particular challenge where lens distortion artifacts can drive large-scale "doming" (Tournadre et al., 2015). While careful camera calibration or the use of drones with built in high precision real time kinematic (RTK) GPS can improve model quality in these cases (Eltner & Sofia, 2020; James & Robson, 2014; Tournadre et al., 2015) the standard way to improve model accuracy is with the addition of **ground control points (GCPs)**.

GCPs are points on the surface of the model with known, externally measured, geographic coordinates. The accuracy, number, and distribution of the ground control points will determine the resulting increase in accuracy of the landscape model (Ferrer-González et al., 2020; Fonstad et al., 2011; Martínez-Carricondo et al., 2018). Across a long and narrow 40 Ha strip similar to what one might film following a group of animals, Ferrer-González et al. (2020) show that when mapping with a quadcopter even three GCPs produce maps with root mean squared error (RMSE) of under .1m in the horizontal direction and under 1m in the vertical direction, but they suggest obtaining more GCPs in order to achieve an error below .03m horizontal RMSE and below .2m in the vertical axis.

GCPs can be generated in a number of ways. In the field, researchers can identify or install physical landmarks that are visible from the air and collect location information for these using a handheld or RTK GPS unit. This approach may be labor intensive for large observation areas, however, and it may be difficult to achieve well-spaced points in landscapes with challenging topography that results in inaccessible areas. Alternatively, if high-resolution georeferenced imagery already exists for the study region, the user can extract location information for landmarks visible in this imagery and assign them to the same landmarks in the landscape model. This approach entails less labor in the field but requires pre-existing imagery of sufficient resolution to allow the user to identify and match an adequate number of landmarks between the two image sources. The quality of the underlying map's georeferencing and the resolution at which GCPs can be matched between the observation model and the existing map will propagate into the final model's accuracy (James et al., 2017).

A compromise solution that we used for our ungulates example is to return to observation sites after the initial observation and conduct systematic drone flights to generate high-quality directly georeferenced landscape imagery. While less accurate than maps generated with GCPs or with RTK GPS-equipped drones, mapping over a larger area with a mapping optimized flight plan can generate maps with internal error less than .5m (Kalacska et al., 2020) and global horizontal RMSE from less than 1m up to nearly 5m depending on the drone used and survey conditions (Eltner & Sofia, 2020; Hugenholtz et al., 2016; James et al., 2017; Kalacska et al., 2020). Particularly, flight plans that include images from multiple heights, crossing flight paths, and oblique camera angles improve model accuracy (Eltner & Sofia, 2020). Such systematic flights can be planned efficiently using many common drone piloting apps, such as Litchi [VC Technology LTD, Brooksville, Florida, USA] and DJI Go 4 [DJI Innovations, Shenzhen, China]. Finally, even including a single GCP improves model absolute accuracy (Eltner & Sofia, 2020). Using this higher quality model, we manually extract coordinates for landmarks visible in both models and use them as GCPs for the observation model.

*4.3 Estimating camera locations for all frames*

We use a simple feature extraction and tracking algorithm to estimate the camera movement, and therefore camera location, between anchor frames. We assume the camera is always pointing towards the ground and can move up and down in relation to the ground and translate or rotate parallel to the ground. For each frame we calculate a partial affine transform that describes how the camera moved to the current frame from a prior reference frame. This movement information is contained in a 2x3 matrix encoding rotation, scaling, and translation. To calculate the transform, we first use the Shi-Tomasi corner detection method in OpenCV (Shi & Tomasi, 1994) and then use the pyramidal Lucas-Kanade method (Bouguet, 1999) to calculate optic flow to estimate the location of these features in the current frame. We use RANSAC (Fischler & Bolles, 1981) to calculate the transform that is most consistent with how the features moved between the frames.

Although this method is typically accurate when there is sufficient overlap between the reference frame and the new frame, the chance of error increases as the overlap between frames decreases. Large errors can occur when there are not enough visual features detected in an image pair either because the overlap between the images becomes too small or the landscape lacks enough distinct features. In this case, the algorithm erroneously calculates movement based on false pairs, leading to arbitrary alignment. To avoid this, one could simply measure the movement between all consecutive frames, since overlap will be high between consecutive pairs, and sum these movements to estimate longer term movement (Haalck et al., 2020; Torney et al., 2018). However, this approach accumulates small errors in movement estimation, which, over many frames, become problematic large scale errors. We compromise by neither registering all frames directly back to the closest anchor frame (risking too few keypoints) nor by registering each frame to its neighbor (risking error accumulation). We instead divide each group of frames between a pair of anchor frames into ten evenly sized subgroups and register each frame to the first frame in its subgroup. The first frame in each subgroup is then registered to the first frame in the subgroup before it, chaining back to the anchor frame at the beginning of the frame group. This increases overlap between frame pairs, decreasing the risk of arbitrary alignment errors, while capping the number of error accumulation steps to ten versus thousands as happens in cases where there are long time gaps between anchor frames. As an added check, we monitor the number of detected features in each image pair and add an additional sub-group beyond the initial ten whenever the number of feature pairs is below fifty.

*4.4 Projecting from pixels in the camera to points on the earth*

Once the landscape topography and camera positions have been defined, we can determine the geographic coordinates for locations in our video frames using the resulting camera or projection matrix, a 3x4 matrix that describes how light passes from the world, through the lens and onto a camera's sensor. Using this information, we can calculate the ray that emanates away from a specific pixel on the camera sensor and out into the world. The place where this ray intersects the ground (or the height of an animal above the ground) in the 3D model of the environment provides an estimate of where the animal is standing on the earth.

To find this ray we write the 3x4 camera matrix P as [M | $p_4$] where M is an invertible 3x3 matrix and $p_4$ is the fourth column of P. x is a given pixel in the video frame (equivalent to a location on the camera sensor) where x = [u, v, 1] and u and v are the height and width from the top left corner of the image frame. This pixel x records anything along the ray defined by

$$X(\mu) = \begin{pmatrix} M^{-1}(\mu \mathbf{x} - \mathbf{p_4}) \\ 1 \end{pmatrix}$$

where $\mu$ is a variable that sets the specific location along the ray (see Hartley & Zisserman, 2004 pp. 161 for a detailed explanation) and $X(\mu)$ is a vector containing [x, y, z] in coordinates of the 3D landscape. We find $\mu$ such that $X(\mu)$ is within a small user-defined threshold away from the ground. For efficient search we start searching along the ray at the last successfully used value of $\mu$ or a fixed starting value if no points have already been found.  If $X(\mu)$ is above the ground we take a step of fixed user defined size farther away from the camera. The process continues until $X(\mu)$ is below ground at which point we take half the sum of the current value of $\mu$ with the last value of $\mu$ where $X(\mu)$ was above ground. We continue in this way taking smaller steps moving back and forth above and below ground until we are within the chosen threshold distance from the ground. The use of the threshold instead of absolute overlap with the ground speeds up the search time. This process is repeated for every location of interest in the observation. For each frame in the observation, we use the affine transform information (see 4.3 Estimating camera locations for all frames) to rotate, scale, and translate the last anchor frame's camera matrix to the current estimated location of the camera before calculating projection rays.

*4.5 Projection validation*

The ultimate georeferenced accuracy of the animal tracks derived from this method depends on how well the landscape model is georeferenced (see 4.2 Georeferencing the landscape model) and how well we can project points from the camera into that landscape. Here we examine the latter. Because we retain the raw video of these observations, this validation process is straightforward. We include a GUI (Fig. S2) to further expedite this process. We choose random individuals and display a frame of the drone video in which they appear. Next to it, we display the general region of the landscape model where we expect that individual to be projected. Looking at the location of the target individual in the raw drone frame, the validator clicks on the equivalent location in the landscape model. We then measure the distance between where the validator clicked and the location our software placed the individual. Doing this many times for different individuals in different frames allows us to estimate the range of expected errors in the observation. We measured median errors of 0.40 m and 0.17 m for the ungulate and gelada worked examples, respectively.

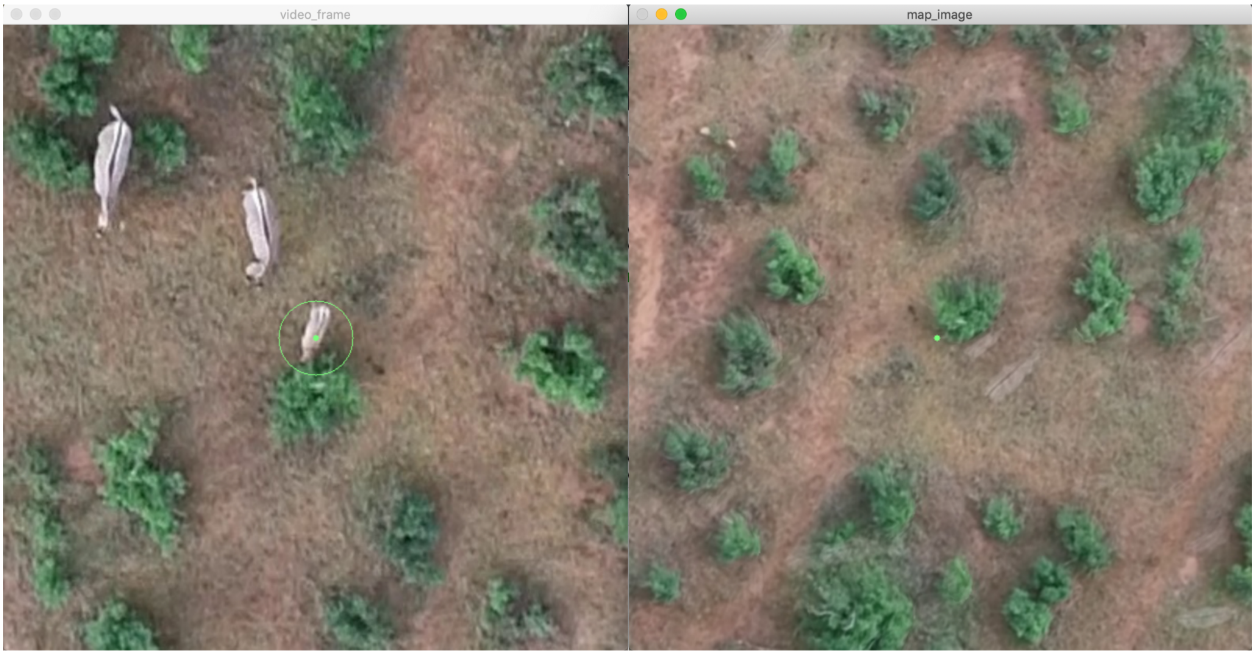


**Figure S2: A visualization of the map projection validation GUI.** On the left is a crop from the drone recording of a randomly chosen individual. On the right is a crop from the SfM-generated landscape orthomosaic around the location where the software expects the individual on the left to be. The validator clicks on the exact location on the image on the right where the zebra is standing. Afterwards, we can compare the distance between the human chosen location and the software derived location. Note, the orientation of the image on the left is determined by the orientation of the drone while filming while the orientation of the image on the left is determined by north (pointing up). Therefore, in this instance, the image on the left is rotated roughly 180 degrees from the image on the right.

*Step 5. Body-part Keypoint Detection*

A key advantage of image-based techniques is that each image can contain much more information than position alone. If animals are filmed at high enough resolution, it is possible to see features like the heads, overall body posture, and in our case with the zebra, even the tails of individuals in the video. Extracting time-varying postural information for individuals can provide a rich dataset for inferring behavior. Inferring behavior is particularly feasible in this context because the low dimensional keypoint information extracted here is paired with video ground truths that the researcher can review any time for verification or annotation.

Automating this process has been revolutionized by easy to use GUI based tools for animal keypoint detection. These tools include SLEAP (Pereira et al., 2020), DeepLabCut (Mathis et al., 2018; Nath et al., 2019), and DeepPoseKit (Graving et al., 2019). Using the animal locations found in Step 3, researchers can extract a small crop, based either on the bounding box size or a fixed area, for each animal in each frame. Using protocols specific to each tool, a subset of these images are annotated and then used to train a model that is applied across all observations. After keypoints have been detected in each crop, those detected positions are put back into the units of the video frame. At this stage, the keypoint positions in the video frame can be transformed into geographic coordinates exactly like the basic locations found in Step 2, as described in Step 4. Additionally, since the locations from Step 2 have already been filtered into corrected trajectories in Step 3, no further tracking at the posture level is necessary. The result is a time-series of locations for each tracked body part for each individual throughout the observation.

In field conditions, animals may become partially or fully occluded by environmental features, such as vegetation, or other animals. Since this keypoint detection step is neural-network-based, like the animal detection step, each keypoint location also has a prediction confidence score. This can be used to filter out low confidence keypoints in frames with occlusions.

Keypoints also allow for finer scale localization of individuals. While the bounding boxes predicted in Step 2 to localize individuals may have some jitter, individual keypoints target multiple specific locations, and can therefore be more stable. Specifically, in the ungulate worked example, we use the mean of the shoulder and hindquarter keypoints to precisely locate the individual in each frame.

It is important to note, however, that the combination of the body shape of the animal being observed, the focal length of the camera lens, and the location of the animal in the video frame can affect the relative geometry of the keypoints for each animal (Fig. S3). This is driven by the fact that animals in the middle of the camera frame will be filmed from directly above while those on the edge of the frame, especially with wider angle lens, will be filmed from a more oblique angle. As a result, the geometry of the keypoints for two individuals standing in the same posture may appear different if one individual stands in the middle of the frame while another stands on the edge. This difference in geometry can make interpreting absolute posture more difficult and can make certain derived metrics like head direction incorrect if frame position is not properly taken into account.

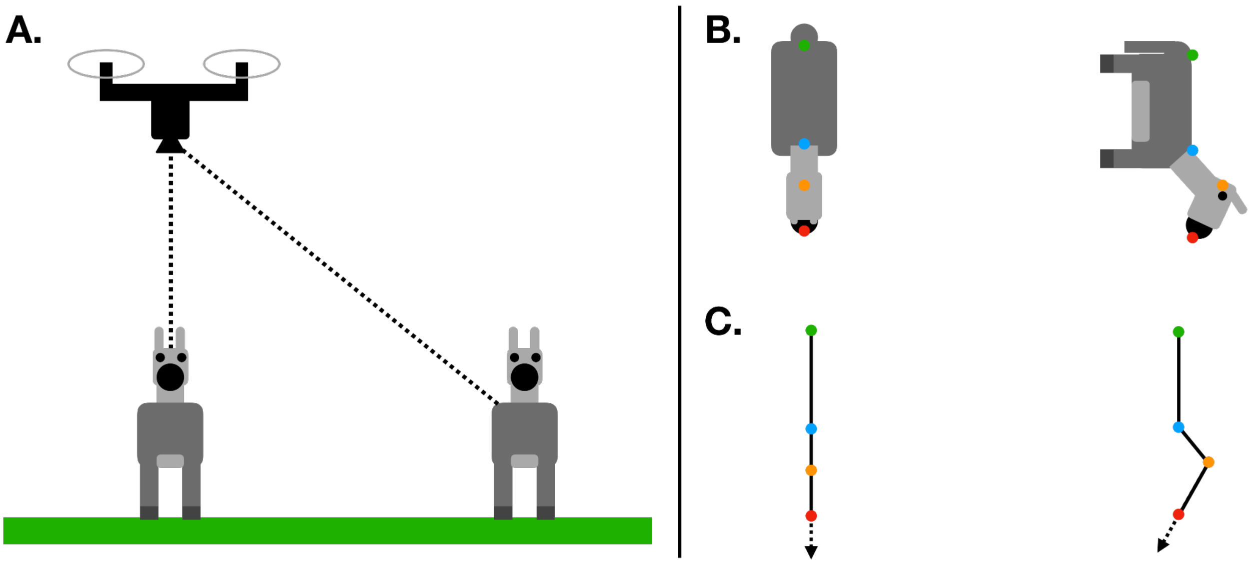


**Figure S3: The effect of an animal's location in the video frame on the relative geometry of its detected body keypoints.** A) When using a drone to film animals from above, animals in the center of the frame will be filmed directly from above, while those on the edge of the frame will be filmed from an oblique angle. B) For animals in identical postures, the difference in filming angle across frame locations leads to different relative geometries among detected keypoints. C) Using the vectors between keypoints to derive behavioral information can give incorrect results if frame position is not accounted for. In this example, even though two animals are facing the same direction with identical body posture, using the vector between the snout (red) and head (orange) keypoints to calculate head direction (dotted arrows) gives different results due to differences in the animals' location within the frame.

*Step 6. Landscape Quantification*

While the primary motivation for building the 3D landscape models in Step 4 is to provide accurate animal tracks without error from drone movement and landscape topography, it also simultaneously provides valuable environmental context for the behavior of the observed individuals. SfM software generates four core outputs which provide information about different aspects of the landscape. The reliability and accuracy of these products come from the choices made while building them in Step 4. These outputs are: 1. An **orthomosaic** which is a single (typically RGB) image of the landscape with camera and landscape distortions removed such that the image has a uniform scale and top-down view throughout (Fig. 5D, E). 2. A **digital surface model (DSM),** a 2D raster image where each pixel encodes the height of the environment in that location including natural and constructed structures, like trees and houses. 3. A **digital terrain model (DTM)**, another 2D raster encoding landscape elevation but excluding surface features like trees and houses. 4. A 3D **point cloud** which is a series of 3D points and associated colors, as recorded by the camera, located on the 3D landscape. The point cloud can be used to generate a 3D **triangular mesh** model (Fig. 5A). Some of these explicitly contain interpretable information about the landscape while others can be further processed to extract biologically-meaningful information.

Researchers can use the DTM and derived slope values to investigate how animals may incorporate landscape topography (which can often be related to energetics) into movement decisions. Similarly, by subtracting the DTM from the DSM, one gets information about the height of above ground objects in the landscape which is directly related to the visual field of each animal within the landscape. In the ungulates worked example we employ a 1.5m threshold above which we consider landscape objects to be a possible visual occlusion. In cases where there is a single dominant surface feature in the landscape, such as a grassland with trees, or a boulder field, one may be able to directly classify large landscape features just based on height. In more heterogeneous environments, however, further processing of the landscape outputs can provide clarity.

Visual color and texture in the orthomosaic (Fig. 5E), either paired with the DSM or alone, provides rich data for computer vision based landscape feature classification. For localized objects, like trees, one can detect and classify them using the same techniques and models as described in Step 2. Many interesting landscape features, such as streams, fences or roads, however, cannot be neatly defined by a bounding box. In those cases, a technique called **semantic segmentation**, in which each pixel in an image is classified as one of a number of classes, is more appropriate. In practice, semantic segmentation is also commonly done with CNNs. Here again, the same core frameworks and general tools as described in Step 2 apply; however, the user will be choosing among a different subset of (pre-built) models. In the ungulates worked example we use semantic segmentation to automatically detect animal trail-like features in the landscape using a model named DeepLabV3 (Chen et al., 2017). Animal trails could be used by animals for, among other reasons, their physical ease of movement or because they record previous animal movements and so may aid navigation (Kashetsky et al., 2021). We were interested in the latter possible case and so used a labeled training set where longer trail-like open ground features were marked as a possible trail but patches of open ground of unclear origin were not considered trails (although they may equally aid physical movement). The output of this approach was a raster map in which each pixel contained the probability, as defined by the trained model, that that pixel overlaps with an animal trail (Fig. 5E, F). The output of our particular trained model is meant to demonstrate the potential of this approach for fine scale landscape quantification but is not intended to be used on its own without more detailed verification.

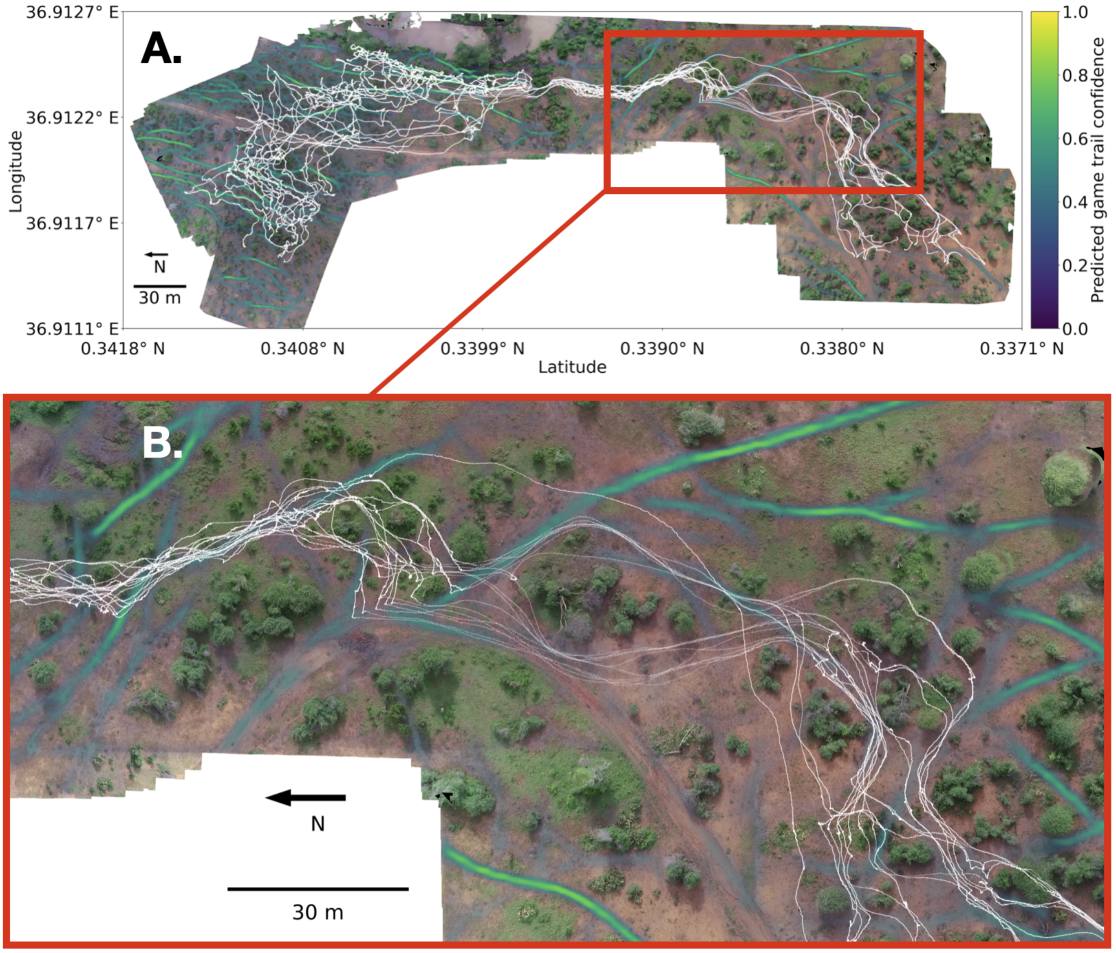


**Figure S4: Predicted animal trails and zebra movement.** A) Using the trained animal trail detection model we overlay the predicted animal trails on the observed environment. Since all the data is in the same coordinate system, we can also overlay the actual movement trajectories of the zebras (white) as they moved through this landscape. B) Zooming in on the part of the observation where the zebras were running from a perceived ground disturbance, we see the effect of the trees and possibly the animal trails on the zebras' movements as they quickly move through the environment.

A challenge with landscape classification of features like animal trails is that unlike the presence or absence of an animal, which is clearly defined (although not always clearly observed), the definition of many functional landscape features vary both with the specific animals that use them and also with the scale at which they are sensed and quantified by researchers (Turner, 1989). Landscape features that may appear consistent and unified in remotely sensed images to a human (i.e. trees) may appear highly variable to the species or even specific animals that encounter them on the ground (Fahrig et al., 2011). Ultimately the specific choice of how a landscape is quantified is a hypothesis presented by the researcher that itself must be tested (Frazier & Kedron, 2017). Supervised deep learning methods for landscape classification have the same limitations since the trained models simply try to approximate human defined labels. The major advantage, however, is that trained models can apply the human driven labels more efficiently since after training no human processing is needed.

A further consideration for using CNN based techniques with orthomosaics is dealing with the large image sizes. Since orthomosaics are often thousands of pixels by thousands of pixels in dimension, they must be split into smaller cropped images before they can be fed into a model. While the exact crop size is bound by the model and hardware chosen for processing, each crop should overlap its neighbors to minimize edge effects during detection or segmentation (Audebert et al., 2017; Y. Liu et al., 2017). In our animal trail example, we use orthomosaic crops of size 2048 pixels x 2048 pixels with an overlap of twenty percent.

**Computing requirements**

Deploying the method described above requires specific computing resources that exceed many personal computers. Computing requirements can be met either with certain high performance gaming-type machines or using dedicated computing clusters including those based in the cloud. Large CNNs need to run on graphical processing units (GPUs) or specialized deep learning processors (for example tensor processing units, TPUs). Specifically, among GPUs, many deep learning frameworks only work with those designed by the NVIDIA [NVIDIA Corporation, USA]. While specific GPU memory requirements depend on the exact model and images, at least 8GB but ideally 10+ GB of memory is generally necessary. SfM mapping tasks, on the other hand, are generally limited by a computer's RAM requiring up to 16GB for mapping projects with 500-1000 images (https://support.pix4d.com/hc/en-us/articles/115002439383-Computer-requirements-PIX4Dmapper). Lastly, storing the high resolution videos and the individual frames that must be extracted from them will require 100s of gigabytes up to terabytes of hard drive space.

| **Term** | **Definition** |
|---|---|
| active learning | An approach related to model-assisted labeling wherein the researcher uses model-generated confidence scores to concentrate annotation effort on difficult examples. |
| anchor frames | The subset of video frames in our method input into the structure from motion software to generate the 3D landscape model and camera location information. |
| annotation | The process of labeling objects of interest in training imagery, for example by drawing bounding boxes around individual animals. |
| bounding box | A rectangle enclosing an object of interest in an image. Bounding boxes can be drawn by the user as a form of annotation, or can be generated by detection models to denote the predicted location of an object of interest. |
| digital surface model (DSM) | A 2D raster image where each pixel encodes the height of the environment in that location, including natural and human-made structures. |
| digital terrain model (DTM) | A 2D raster image encoding landscape elevation, excluding surface structures such as trees and buildings. |
| graphical user interface (GUI) | A means of viewing or inputting data that relies on graphical elements (e.g. buttons) rather than coding inputs. |
| ground control point (GCP) | Locations or landmarks with known real-world geographic coordinates. These points are used to georeference the landscape model generated from the anchor frames. |
| image augmentation | A technique for increasing the effective size of the training set by modifying each training image each time it is shown to the model. Modifications, including blurring, rotating, and adjusting contrast, and are intended to mimic the diversity of images in the entire dataset. |
| model-assisted labeling | An iterative process where image annotation and model training are conducted in parallel. The user initially labels a small number of images and uses these to train an initial version of the model. This initial model is then used to generate annotations for the remaining images, which the user then confirms or corrects while also adding annotations for any animals that the model missed. |
| orthomosaic | Two-dimensional composite images generated by the SfM software that approximate the appearance of 3D structures from an overhead |

| | |
|---|---|
| | viewpoint. |
| point cloud | A set of 3D points often also containing a color value that can be used to represent landscapes or objects in space. |
| pretraining | A feature of many common deep learning models where the model has been initially trained on large datasets consisting of generic imagery of common scenes and objects. This allows the model to learn the basic, universal aspects of imagery before being fine-tuned on the user's specific dataset. |
| semantic segmentation | The process of labeling every pixel in an image as one of a set number of object classes. |
| structure from motion (SfM) | A technique for 3D reconstruction in which two-dimensional images from various overlapping viewpoints are used to define the geometry of the target structure (here, the landscape surrounding the observed animals). |
| triangular mesh | A 3D surface model created from a point cloud by connecting triads of points to create flat triangular surfaces. |

**Table S2. Glossary of terms.** Defined terms are bolded at first appearance in the supplement.
